# Periplasmic production of Green Fluorescent Protein is poorly tolerated by *Escherichia coli*

**DOI:** 10.1101/2025.09.28.679025

**Authors:** Alexander Osgerby, Tim W Overton

## Abstract

*Escherichia coli* is a commonly used host for recombinant protein production. It is advantageous to direct many recombinant proteins, especially those that require disulphide bonding for function, such as antibody fragments, to the periplasm of *E. coli*. This requires N-terminal fusion of a signal peptide that directs the polypeptide chain through the relevant translocation apparatus. Signal peptides cannot be selected on the basis of recombinant gene sequence, so screening is required to select the optimal signal peptide for each product, typically using subcellular fractionation, a time-intensive process. Fusion of a fluorescent protein such as GFP to the C-terminal of recombinant proteins has previously been used to accelerate cytoplasmic protein production process development, but most GFP proteins are not active in the periplasm. Previous studies have developed GFP derivatives that fold rapidly (such as superfolder GFP, sfGFP) and have been reported to be periplasmically active. Here, we tested the applicability of sfGFP as a periplasmic screening tool using single-cell analysis and structured illumination microscopy. We discovered that sfGFP is very poorly tolerated in the periplasm, causing deleterious effects on *E. coli* physiology, manifesting as poor growth, cell death, and loss of recombinant protein productivity. A further reason for poor GFP functionality in the periplasm is errant disulphide bonding, so we tested a cysteine-free GFP, which cannot form disulphide bonds; results were similar to sfGFP. In conclusion, currently-available GFP variants are poor fusion partners for screening production and translocation of recombinant proteins to the *E. coli* periplasm due to their negative impact on physiology.

**Highlights:**

- GFP is a useful screening tool recombinant protein production.
- We tested periplasmic expression of GFP derivatives sfGFP and cfSGFP2.
- We used structured illumination microscopy to visualise GFP accumulation.
- Periplasmic GFP derivatives have significant negative effects on bacterial physiology

## Introduction

Recombinant protein production (RPP) is an industrially- and academically-important technology with applications across multiple sectors. The bacterium *Escherichia coli* is an important host for RPP [1]. In the biopharmaceutical industry, *E. coli* is used to manufacture many smaller, relatively simple proteins that do not require extensive post-translational modification such as most insulin products, human growth hormone, and colony-stimulating factor [2].

Rapid optimisation of RPP is a key step in development of new processes, enabling high yields of recombinant protein. A relatively common high-throughput optimisation method is translational fusion of an autofluorescent protein such as green fluorescent protein (GFP) to the C-terminus of the RP of interest. Measurement of green fluorescence enables detection of not only the quantity of RP-GFP fusion but also determination of its folding state; typically, correctly-folded RP results in functional GFP with bright fluorescence, whereas incorrectly-folded RP decreases GFP fluorescence [3–5]. Previous work has demonstrated that production of GFP in the *E. coli* cytoplasm is well-tolerated; GFP can accumulate to high levels and does not massively negatively impact growth and viability, unlike some other RPs which can cause plasmid loss and cell death [6–9].

The periplasmic space, located between the inner and outer membranes of Gram-negative bacteria, is an advantageous compartment to localise certain RPs to, especially those requiring disulphide bonding, for example antibody fragments [10,11]. Not only is the periplasm oxidizing, but it is also the location of the Dsb enzymes which catalyse disulphide bond formation and isomerisation [12]. Periplasmic localisation also permits simpler methods for release of RP than the cytoplasm [13].

Proteins (both recombinant and native) are translocated across the inner membrane via one of three pathways in *E. coli*. The Tat apparatus exports fully-folded proteins [14]; there are presently 27 known native Tat substrates in *E. coli* [15]. The Sec apparatus is the major pathway for translocation in *E. coli* [16]. Sec translocates unfolded polypeptide chains via one of two routes: the post-translational route, mediated by SecA, whereby fully-translated polypeptides are translocated through the SecYEG complex [17]; and the co-translational route, mediated by SRP, where translocation of a polypeptide chain proceeds while it is still emerging from the ribosome [18]. In each case, an N-terminal signal peptide directs the protein / polypeptide to the required translocase [19]. There is currently no way to predict which signal peptide will be best for each recombinant protein, so screening is commonly required [19,20]. There are few methods for high-throughput quantification of periplasmic accumulation and folding of RPs in *E. coli* (reviewed by [21]) so many screening approaches require time-consuming subcellular fractionation followed by gel electrophoresis and Western blotting.

Periplasmic localisation poses a problem for most GFP variants [22], therefore GFP has not been widely used for screening. Once translocated through Sec in an unfolded form, the two cysteine residues in GFP (C48 & C70) become aberrantly disulphide bonded in the oxidizing periplasm (either to each other or to other proteins), resulting in GFP misfolding and lack of fluorochrome maturation [22,23]. GFP can be successfully translocated by Tat [24], although Tat is not as widely used as Sec in RPP.

The first Sec-compatible GFP reported was superfolder GFP (sfGFP), developed as an enhanced-folding GFP derivative [25]. Directed evolution was used to generate sfGFP from folding reporter GFP [26] which contains the three “cycle-3” mutations (F99S, M153T, & V163A; [27]) and the two eGFP mutations (F64L & S65T) derived from GFPmut1 [28]. The resultant evolved sfGFP contained an addition six mutations (S30R, Y39N, N105T, Y145F, I171V & A206V) giving rise to improved folding kinetics.

Dinh & Berhhardt [29] directed sfGFP to the *E. coli* periplasm using three approaches, using fluorescence microscopy and subcellular fractionation and immunoblotting to confirm localisation. Fusion to the *malE* gene (encoding maltose binding protein) under direction of the *malE* signal peptide (MalE^sp^, targeting the post-translational Sec pathway) resulted in periplasmic localisation, with accumulation at cell poles. Fusion to the *malE* signal peptide (MalE^sp^) did not result in significant periplasmic accumulation. However, fusion to the *dsbA* signal peptide (DsbA^sp^), reportedly targeting via the co-translational branch of Sec, resulted in periplasmic localisation. DsbA^sp^ was initially described as a co-translational signal peptide [30] but has more recently been described as being unlikely to be an SRP substrate [31].

These findings were noted to be different to those of Fisher & DeLisa [32], who found that DsbA^sp^ directed only a small proportion of sfGFP to the periplasm, where it was inactive. This discrepancy was proposed to be due to expression levels; Dinh & Bernhardt used lower expression from a chromosomal insertion under control of the *lac* promoter, whereas Fisher & DeLisa utilised higher expression (a multi-copy plasmid under the stronger *trc* promoter). This suggests that successful periplasmic translocation of sfGFP could be highly dependent upon expression strength.

Finally, Aronson *et al*. [33] described periplasmic localisation of sfGFP when translocated by the MBP^*^1 signal peptide, a mutated variant of MalE^sp^ that is more hydrophobic and is reported to direct translocation via the co-translational pathway [34].

Previous studies did not report the effect of periplasmic sfGFP accumulation on bacterial physiology. RPP can induce stress in *E. coli* through a variety of pathways, for example metabolic burden caused by division of metabolites and energy between RPP and biomass production, stringent response, and RP misfolding leading to a heat shock response [1,35,36]. Periplasmic RPP potentially adds additional stresses caused by bottlenecks at Sec, accumulation of untranslocated (and potentially misfolded) polypeptides in the cytoplasm, and misfolding of translocated polypeptides in the periplasm.

In this study we sought to understand the utility of GFP derivatives as a screening tool for measuring the quantity and folding quality of RP in the *E. coli* periplasm. A key objective was understanding the quantity of GFP that could be successfully translocated into the periplasm and its impact on bacterial physiology. Periplasmic RPP requires accumulation of relatively large quantities of RP in the periplasm, which could negatively impact on physiology. As such, a screening method using GFP must not in and of itself negatively affect bacterial physiology.

Following determining the utility of sfGFP for screening, including using super-resolution structured illumination microscopy to visualise sfGFP accumulation, we used a cysteine-free GFP, cfSGFP2, for the first time in *E. coli*. Our results suggest that different GFP derivatives are of limited use for screening in the bacterial periplasm due to significant negative effects on bacterial physiology, manifesting as loss of growth, decreases in viability, and reduced accumulation of recombinant protein.

## Materials and Methods

### Strains and plasmids

*E. coli* strains BL21 (F^−^ *ompT gal dcm lon hsdS*_B_(r_B_^−^ m_B_^−^) [*malB*^+^]_K-12_(λ^S^); [37]) and BL21(DE3) (BL21 λ(DE3 [*lacI lacUV5*-T7p07 *ind1 sam7 nin5*]) [37]) were used for expression studies. Plasmid construction was performed using *E. coli* DH5α *fhuA2* (*fhuA2Δ*(*argF-lacZ*)U169 *phoA glnV44* Φ80Δ(*lacZ*)M15 *gyrA96 recA1 relA1 endA1 thi-1 hsdR17*; New England Biosciences, [38]). Plasmids are listed in Supplementary Table S1. DNA primers are listed in Supplementary Table S2. Full details of plasmid construction are included in the supplementary information.

### Growth conditions

Cultures were grown in 250 mL conical flasks in a MaxQ 4000 incubator (Thermo Scientific) at 30 °C and 200 rpm. Cultures of BL21(DE3) were grown in 50 mL of Terrific Broth (TB: 20 g·L^-1^ Tryptone; 24 g·L^-1^ Yeast Extract; 17 mM KH_2_PO_4_; 72 mM K_2_HPO_4_; 0.4% (v/v) glycerol, pH 7) with 0.4% (w/v) glucose. Cultures of BL21 were grown in 50 mL of TB. Kanamycin sulphate (Sigma) was added at 50 µg·mL^-1^ for growth of transformants. Isopropyl β-D-1-thiogalactopyranoside (IPTG; dioxane free, Anatrace) and Arabinose (Thermo) were used as inducers. Optical density was measured using an Evolution 300 UV-Vis spectrophotometer (Thermo Scientific) at 600 nm, in disposable plastic cuvettes.

### Fractionation and SDS-PAGE

Periplasmic and cytoplasmic fractions were separated according to [20]. Samples were thawed on ice, mixed 1:1 with 15 µL 2X SDS PAGE sample buffer (65.8 mM Tris-HCl pH 6.8, 0.01% bromophenol blue, 26.3% glycerol, 2.1% SDS) and 1.5 µL of 1 M dithiothreitol (DTT), then heated at 95°C for 5 min. Samples were separated by SDS-PAGE using Novex™ WedgeWell™ 16%, Tris-Glycine, 1.0 mm, 12-well, Mini Protein Gels (Thermo Scientific) at 140 V for 45 min using a mini gel tank (Life Technologies). 30 µL of each sample was loaded and 5 µL PageRuler Prestained protein ladder (Thermo Scientific) was run in parallel. Gels were stained using SimplyBlue SafeStain (Invitrogen) according to manufacturer’s instructions and imaged.

### Flow cytometry

Samples were analysed using a BD Accuri C6 (Fig 1,4B-C)) or BD Accuri C6 Plus (Fig 4E-F, 7) flow cytometer (Becton Dickinson). Samples were diluted in 0.2 µm filter sterilised Phosphate Buffered Saline (PBS). Dead cells were detected using propidium iodide (PI, final concentration of 4 µg·mL^-1^, incubated for 5 min in the dark. A flow rate of 14 µL·min^-1^, core size of 10 µm and primary threshold of 12,000 on FSC-H were used with an event rate < 4000 events·s^-1^. 25,000 events were measured per sample. Cells were excited by a 488 nm laser and fluorescence detected using 533/30 BP (GFP) and 670 LP (PI) filters. Data were analysed using BD Accuri C6 or BD Accuri C6 Plus software (Becton Dickinson).

**Figure 1.**
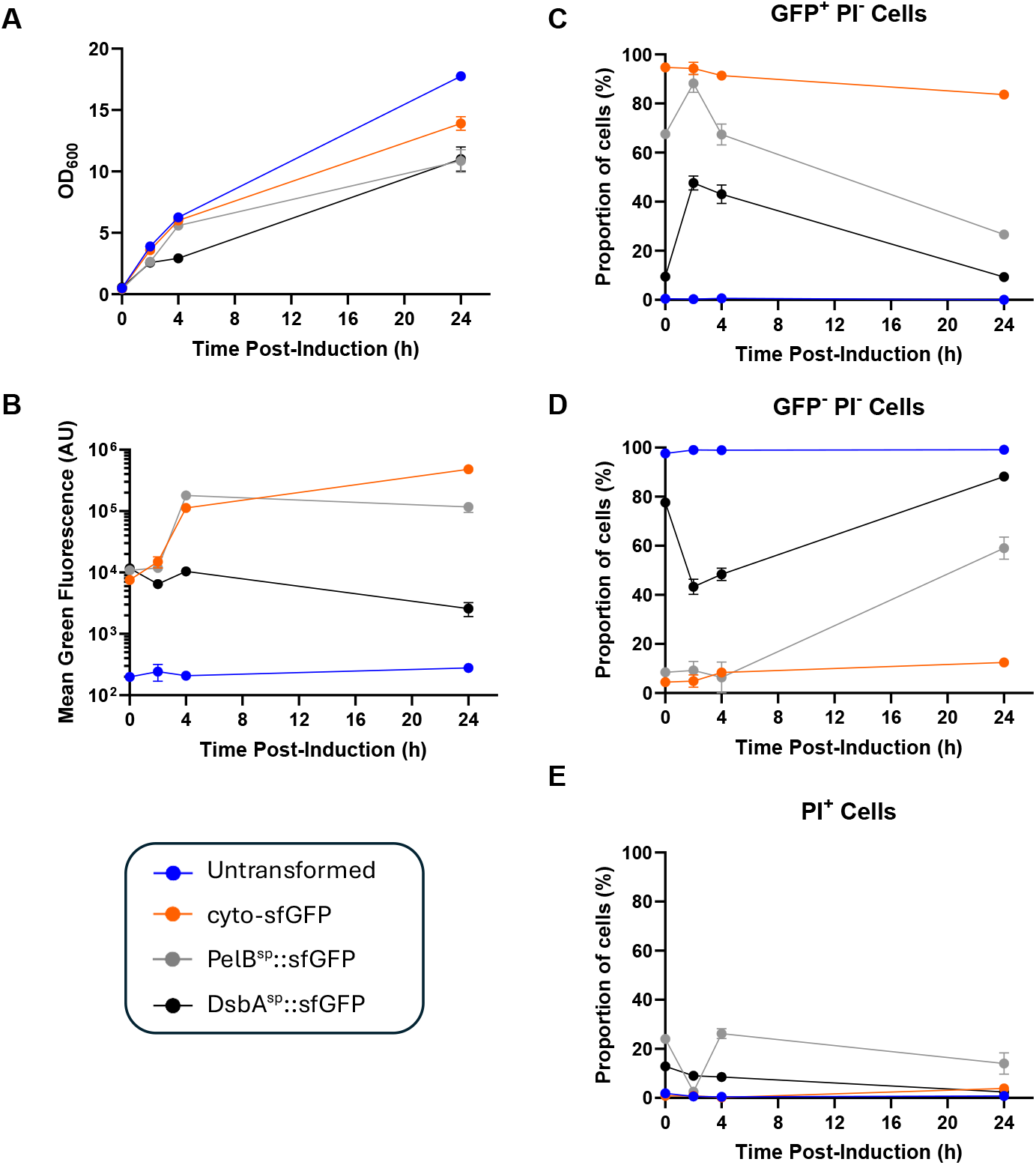
Expression of sfGFP from the *T7lac* promoter. *E. coli* BL21(DE3) transformed with either pET-26b(+)Δ-ATG-msfgfp (expressing cytoplasmic sfGFP), pET-26b(+)-PelB^SP^::msfgfp (periplasmic targeting using PelB^sp^) or pET-26b(+)Δ-DsbA^SP^::msfgfp (periplasmic targeting using DsbA^sp^) were grown in terrific broth at 30 °C with shaking at 200 rpm until an OD_600_ ∼ 0.4-0.6, whereupon 50 μM IPTG was added. Samples were taken at regular intervals and OD_600_ was measured (**A**). Flow cytometry was used to measure green fluorescence (**B**), and the percentage of cells that were alive (PI^-^) and productive (GFP^+^, **C**), alive and non-productive (GFP^-^ PI^-^, **D**), or dead (PI^+^, **E**). Error bars represent +/- SD from duplicate flasks.

### Fluorescence Microscopy

Cells expressing GFP were imaged using a Zeiss Axiolab microscope and Moticam 10+ camera. Cells were diluted to OD_600_ 0.1 in sterile diH_2_O, 2 µL transferred to 1 mm glass slides (Fisher) topped with #1.5 glass cover slips (Fisher). Immersion oil (PanScan Xtra) with a refractive index of 1.518 was added just prior to switching to the 100x oil immersion objective (Zeiss).

### Structured Illumination Microscopy

Bacteria were imaged by structured illumination microscopy (SIM) using a Nikon N-SIM S. Cells were induced with IPTG and samples taken after 2 h and diluted to OD_600_ 0.4 in sterile diH_2_O. 30 µM Nile red was added for 15 min, centrifuged at 5000 x g for 5 min, and the pellet resuspended in sterile diH_2_O. For imaging, 2 µL of cell resuspension was transferred to agarose pads prepared according to [39] using 1 % low-melt agarose (VWR) dissolved in diH_2_O and topped with #1.5 glass cover slips (Fisher).

Immersion oil (Nikon) with a refractive index of 1.518 was added just prior to imaging using a 100x oil immersion objective. GFP was excited using a 488 nm laser and emission detected using a Chroma ET525/50m bandpass filter. Nile red was excited with a 561 nm laser and emission detected using a Chroma ET 700/75m bandpass filter. Images were acquired using Nikon NIS Elements 5 and analysed using NIS Elements 5 workstation, and processed using ImageJ [40].

## Results & Discussion

### Periplasmic production of sfGFP from a pET vector

Our starting point was translocation of sfGFP to the periplasm. We chose two widely-used signal peptides: the *pelB* signal peptide (PelB^sp^) from *Erwinia carotovora*, which directs to the post-translational export pathway; and the *E. coli dsbA* signal peptide (DsbA^sp^). We have previously shown that PelB^sp^ and DsbA^sp^ have very different functionalities when used to direct periplasmic translocation of an scFv antibody fragment [20], influencing not only translocation, but also the level of RP accumulation and bacterial physiology. Expression of sfGFP was driven from T7*lac* promoter on the pET-26b vector. Three constructs were generated, expressing sfGFP with PelB^sp^, DsbA^sp^, or no signal peptide and thereby directing cytoplasmic expression (referred to as cyto-sfGFP).

BL21(DE3) transformed with these plasmids (and an untransformed control) were grown in terrific broth in shakeflasks at 30 °C until OD_600_ of 0.4-0.6 whereupon they were induced with 50 μM IPTG. Comparison of growth curves (Fig. 1A) show that there was little growth inhibition in cultures producing cyto-sfGFP, except for a lower final yield than untransformed cultures. PelB^sp^ induced a growth defect 2 h after induction and a decreased final biomass yield, whereas DsbA^sp^ resulted in a more severe growth defect. It should also be noted that PelB^sp^::sfGFP and DsbA^sp^::sfGFP transformants grew far more slowly than cyto-sfGFP and untransformed cells; the quantity of inoculum had to be increased to permit growth to the required biomass concentration within a reasonable timeframe. Flow cytometry analysis of green fluorescence (Fig. 1B) showed high fluorescence in cyto- and PelB^sp^::sfGFP cultures but far lower fluorescence for DsbA^sp^. It should be noted that green fluorescence of all transformed cultures was high before induction, due to the inherent leakiness of pET systems. Indeed, DsbA^sp^ cultures did not increase in green fluorescence after induction. Inspection of green fluorescence histograms (Supplemental Figure S1) reveals single populations for untransformed cultures through growth; cyto-sfGFP and PelB^sp^::sfGFP cultures started as single populations and become more heterogenous at 24 h likely caused by plasmid loss (the lowest fluorescence sub-population being equivalent to plasmid free cells). DsbA^sp^::sfGFP cultures were heterogenous throughout; at induction, comprising three populations (the lowest having comparable fluorescence to untransformed cells and therefore likely to be plasmid-free), following induction having two populations (GFP^-^ and GFP^+^), and mainly comprising the GFP^-^ population at 24h.

Flow cytometry with propidium iodide staining (PI stains dead cells) was also used to differentiate between live GFP^+^, live GFP^-^ and dead cells (Fig. 1C-E). Whereas untransfomed and cyto-sfGFP cultures had low proportions of dead (PI^+^) cells throughout growth, periplasmic sfGFP cultures had far higher levels of cell death, especially for PelB^sp^. The proportion of live, productive (GFP^+^) cells was highest for cyto-sfGFP, then PelB^sp^, then DsbA^sp^, suggesting that periplasmic targeting resulted in loss of productivity and cell death.

Samples from cultures were fractionated to separate periplasmic and cytoplasmic proteins and analysed by SDS-PAGE (Fig. 2). In cultures expressing cyto-sfGFP, a protein band of ∼ 27 kDa in the cytoplasmic fraction increased in intensity following induction. However, a band of a similar size was also visible in the periplasmic fraction; for the 2 h and 4 h samples, there was greater intensity in the cytoplasmic fraction, whereas at 24 h the periplasmic fraction band intensity appeared higher. This is likely a consequence of the negative effect of physiology leading to a loss in envelope integrity; subcellular fractionation (which relies upon a robust, intact envelope) could not precisely separate cytoplasmic and periplasmic fractions, and some cytoplasmic proteins (including sfGFP) passed into the periplasmic fraction.

**Figure 2.**
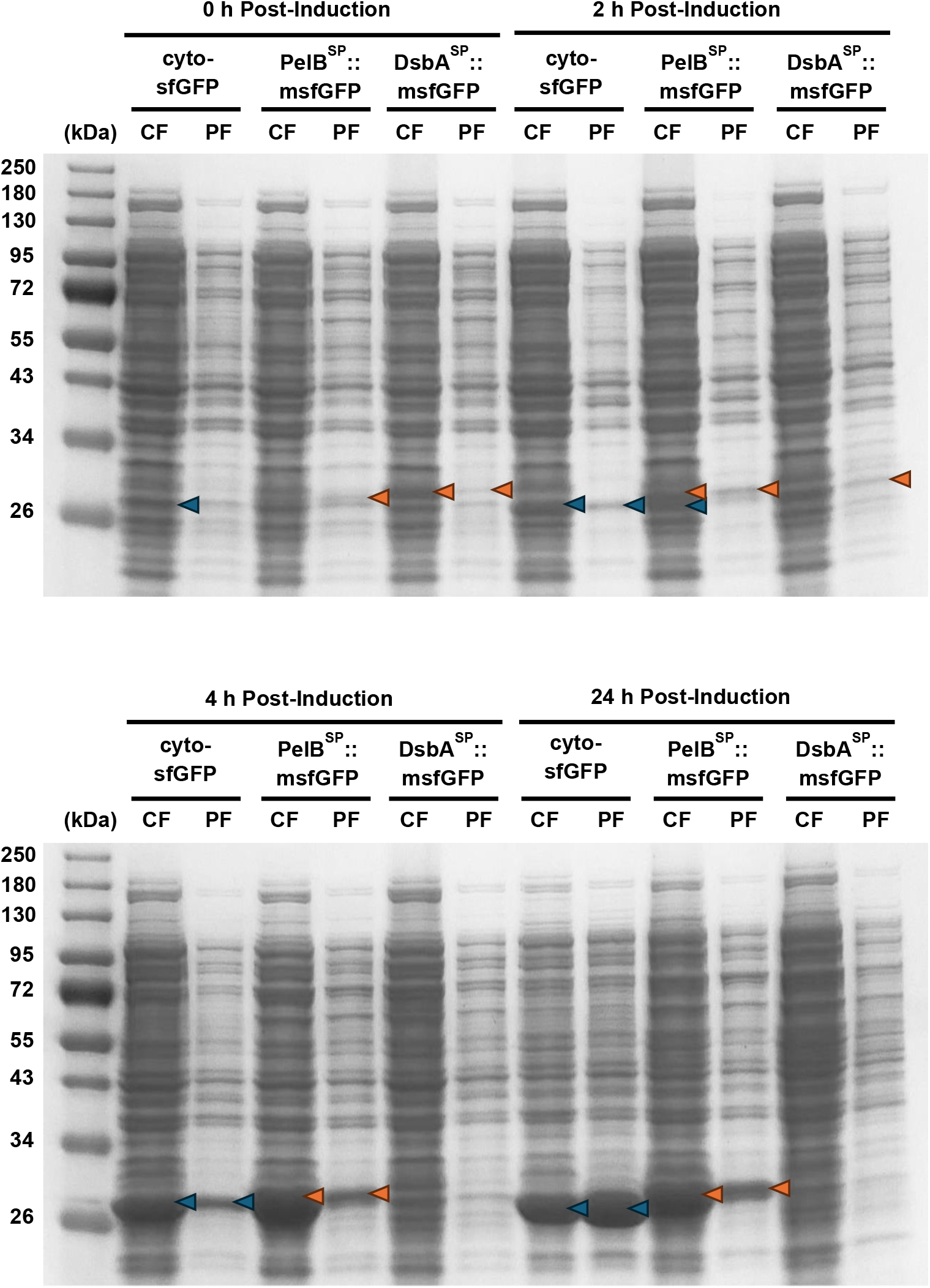
SDS-PAGE analysis of sfGFP expressed from the *T7lac* promoter. *E. coli* BL21(DE3) transformed with either pET-26b(+)Δ-ATG-msfgfp (expressing cytoplasmic sfGFP), pET-26b(+)-PelB^SP^::msfgfp (periplasmic targeting using PelB^sp^) or pET-26b(+)Δ-DsbA^SP^::msfgfp (periplasmic targeting using DsbA^sp^) were grown in terrific broth at 30 °C with shaking at 200 rpm until an OD_600_ ∼ 0.4-0.6, whereupon 50 μM IPTG was added. Samples were taken at regular intervals, fractionated into cytoplasmic and periplasmic fractions, and separated by SDS-PAGE. Bands corresponding to sfGFP are indicated by arrows, blue referring to sfGFP and red to sfGFP fused to a signal peptide with a resultant higher molecular weight.

In cultures expressing PelB^sp^::sfGFP, corresponding bands appeared in the periplasmic fractions after 2 h which appeared to be of a higher molecular weight than that observed in cyto-sfGFP cultures, likely sfGFP with an intact PelB^sp^, denoting inefficient cleavage by signal peptidase. The resolution of the SDS-PAGE gel was not sufficient to discriminate between the uncleaved PelB^sp^::sfGFP and the cleaved variant, so the efficiency of cleavage cannot be determined. The inefficiency of subcellular fractionation (likely due to the negative effects on physiology leading the envelope fragility) means that accurate subcellular localisation was not possible. Fluorescence microscopy (Fig. 3A) revealed that some bacteria were uniformly green and some had punctate green fluorescence (potentially inclusion bodies); no “halo” of green fluorescence, previously suggested to indicate periplasmic localisation [29,33], was observed. Taken together, PelB^sp^::sfGFP appeared to accumulate in a comparable manner to cyto-sfGFP (corroborating FCM fluorescence data, Fig. 1B) but not successfully localise to the periplasm; stress caused by PelB^sp^::sfGFP synthesis resulted in severe growth inhibition, cell death and loss of productivity (Fig. 1B-E).

**Figure 3.**
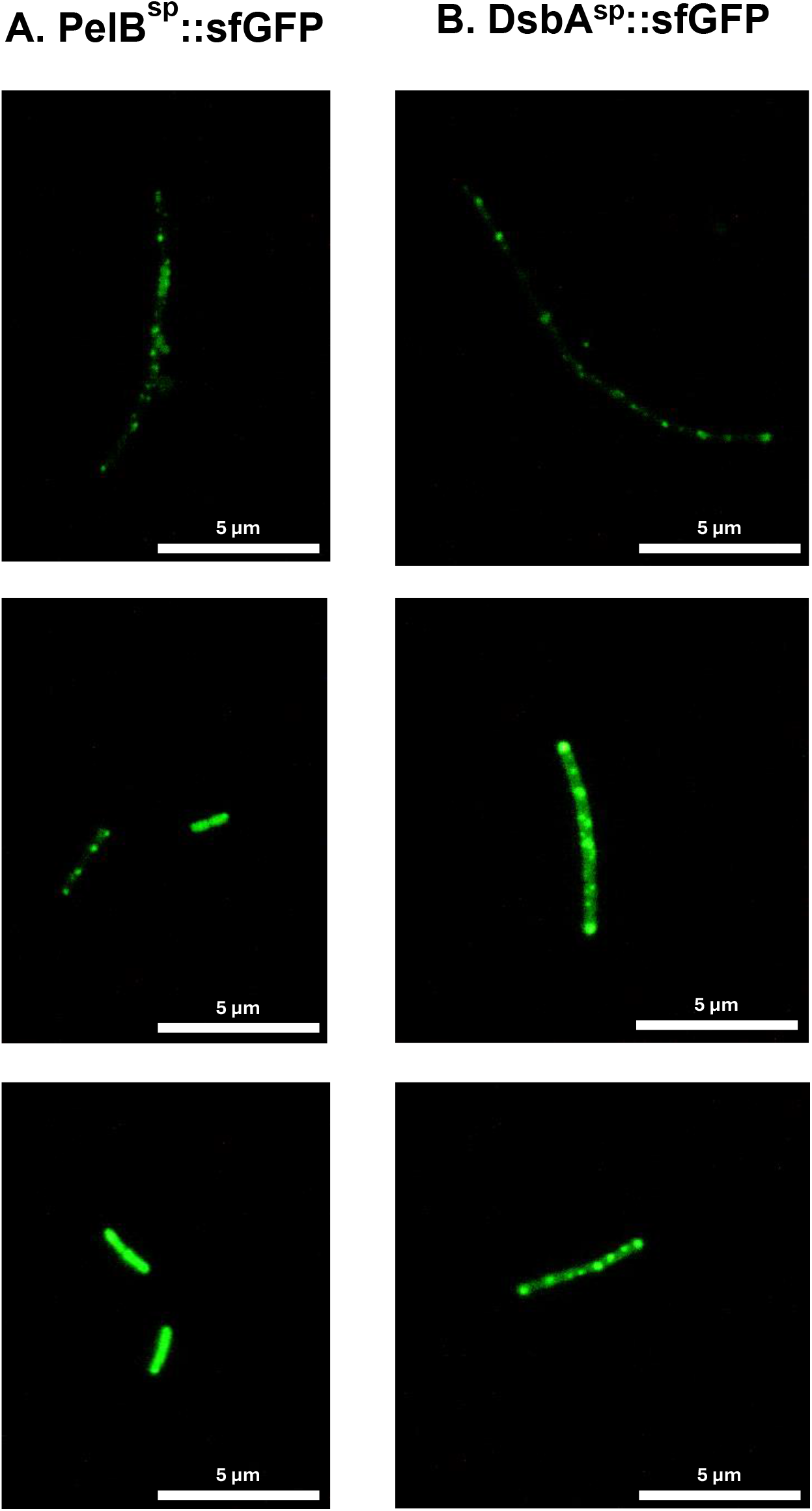
Fluorescence microscopy of sfGFP expressed from the *T7lac* promoter. *E. coli* BL21(DE3) transformed with either pET-26b(+)Δ-ATG-msfgfp (expressing cytoplasmic sfGFP), pET-26b(+)-PelB^SP^::msfgfp (periplasmic targeting using PelB^sp^) or pET-26b(+)Δ-DsbA^SP^::msfgfp (periplasmic targeting using DsbA^sp^) were grown in terrific broth at 30 °C with shaking at 200 rpm until an OD_600_ ∼ 0.4-0.6, whereupon 50 μM IPTG was added. At 4 h post-induction cells were visualised using fluorescence microscopy. Error bars represent +/- SD from duplicate flasks.

SDS-PAGE analysis (Fig. 2) revealed far lower levels of DsbA^sp^::sfGFP accumulation than cyto-sfGFP or PelB^sp^::sfGFP, corresponding to fluorescence data (Fig. 1B). Fluorescence microscopy (Fig. 3B) showed punctate green fluorescence, as previously reported [33], although no “halo”. In summary, use of DsbA^sp^ resulted in far lower sfGFP productivity but still had a detrimental effect on physiology, suggesting that stress was mainly generated by protein translocation and folding rather than metabolic burden. We had previously observed that use of DsbA^sp^ to target an scFv to the periplasm had a more detrimental effect of physiology than PelB^sp^ [20], although scFv accumulation per unit biomass was higher for DsbA^sp^.

Overall, sfGFP is not amenable to screening of periplasmic protein production when regulated by the *T7lac* promoter as bacterial physiology is significantly negatively affected. We therefore switched to a more tightly-regulated promoter, the arabinose-inducible pBAD system [41].

### Use of a pBAD expression vector

Expression constructs based on pBAD30 [41] were constructed with either no signal peptide, PelB^sp^, or DsbA^sp^; the native ampicillin resistance of pBAD30 was replaced with kanamycin resistance for tighter selection. pBAD30 also was chosen as it has a p15A origin of replication and thereby a lower copy number than the pBR322 origin of pET26b, reducing gene dosage.

Unlike the *T7lac* promoter system, overnight cultures containing pBAD constructs had very low green fluorescence, therefore little sfGFP expression, demonstrating lack of leakiness. Induction of sfGFP production with 0.2 % arabinose resulted in similar growth inhibition as the *T7lac* promoter cultures for cyto-sfGFP and DsbA^sp^::sfGFP, but greater inhibition for PelB^sp^::sfGFP (Fig. 4A). Green fluorescence of cyto-sfGFP cultures was highest, followed by PelB^sp^::sfGFP then DsbA^sp^::sfGFP; all cultures showed a decrease in fluorescence from 4 h to 24 h. Green fluorescence histograms (Supplemental Fig. S2) showed low levels of culture heterogeneity, with small numbers of non-fluorescent cells at 2 and 4 hours (∼10^2^ fluorescence units); DsbA^sp^::sfGFP cultures were mainly non-fluorescent at 24 h, suggesting plasmid loss as a result of physiological stress. The proportion of dead cells (Fig. 4C) was far higher for periplasmic sfGFP, comparable to the T7lac promoter system (Fig. 1E).

**Figure 4.**
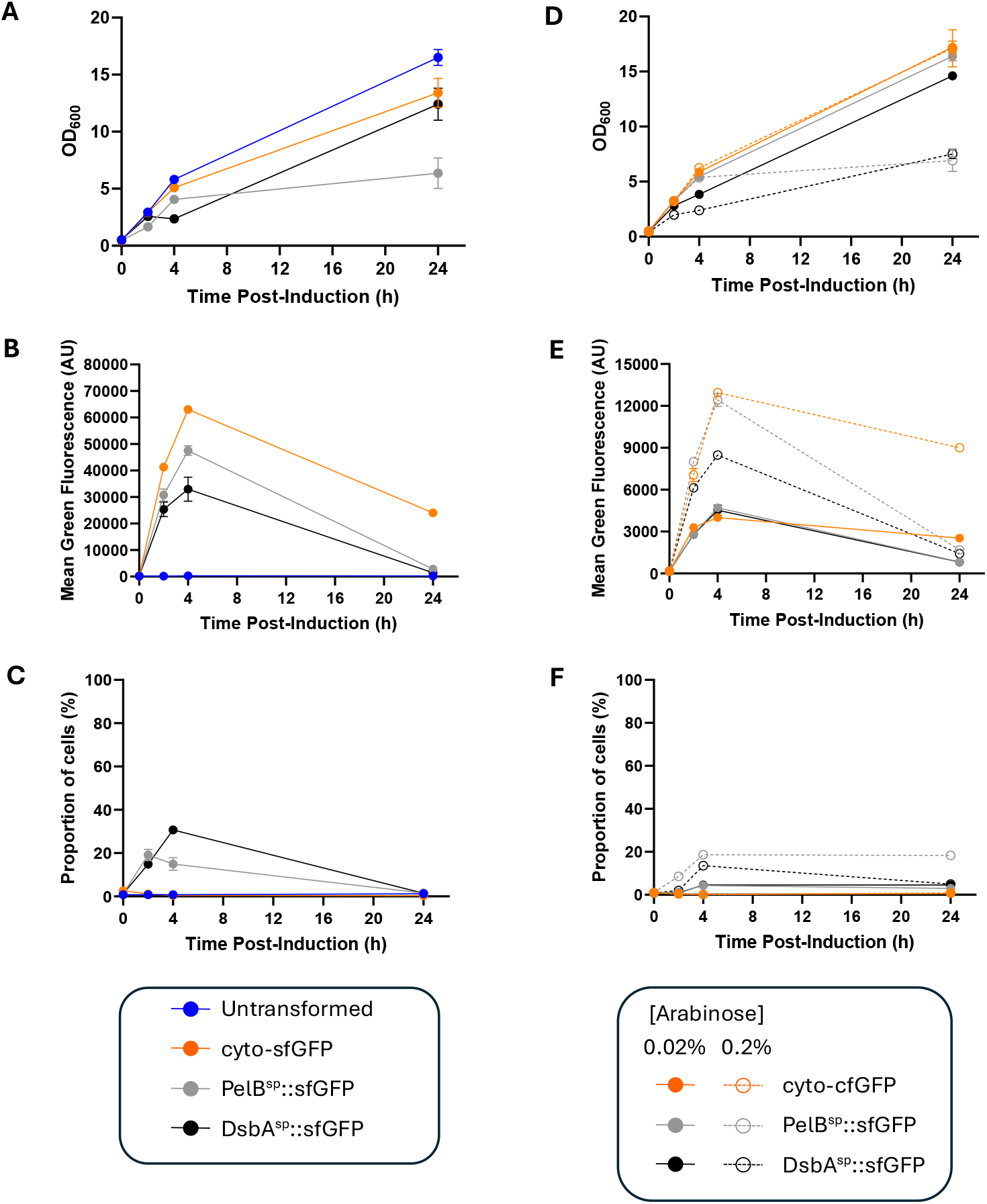
Expression of sfGFP from the *pBAD* promoter. *E. coli* BL21 transformed with either pBAD30-kan^R^-pelB^SP^::msfgfp, pBAD30-kan^R^-dsbA^SP^::msfgfp, or pBAD30-kan^R^-msfgfp were grown in terrific broth at 30 °C with shaking at 200 rpm until an OD_600_ ∼ 0.4-0.6, whereupon 0.2 % (**A-F**) or 0.02 % (**D-F**) arabinose was added to induce RPP. Samples were taken at regular intervals and OD_600_ was measured (**A**,**D**). Flow cytometry was used to measure green fluorescence (**B**,**E**), and the percentage of cells that were dead (PI^+^, **C**,**F**). Note that a different flow cytometer was used for each experiment (panels D&C / E&F), hence the absolute green fluorescence values are not comparable. Error bars represent +/- SD from duplicate flasks.

Given that 0.2 % arabinose still elicited growth arrest and cell death, the experiment was repeated using induction with 0.2 % or 0.02 % arabinose. The lower arabinose concentration resulted in far less growth arrest, with PelB^sp^::sfGFP growing in a similar manner to cyto-sfGFP, although DsbA^sp^::sfGFP was still growth inhibited (Fig. 4D). Mean green fluorescence was comparable for periplasmic and cyto-sfGFP with 0.02 % arabinose induction (Fig. 4E) although periplasmic fluorescence declined at 24 h. Single-cell analysis shows that 0.02 % arabinose cultures are heterogenous at 2 h and 4 h, indicative of partial induction (Supplemental Fig. S3), and 24 h periplasmic cultures are largely non-fluorescent. Cell death was far lower at 0.02 % arabinose than 0.2 % (Fig. 4F). It can be seen that one must balance higher protein production but poor growth at 0.2 % arabinose against lower protein production, culture heterogeneity and only partial induction, with better growth at 0.02 %. This means that screening would need to be performed using relatively low expression levels; subsequent intensification of growth (eg fed-batch bioreactors, higher induction) would likely result in poor growth and physiology, severely limiting this as a screening tool.

SDS-PAGE analysis (Supplemental Fig. S4) confirmed higher accumulation of cyto-sfGFP and PelB^sp^::sfGFP than DsbA^sp^::sfGFP, higher accumulation in 0.2 % arabinose cultures than 0.02 % cultures, and a decrease in sfGFP in periplasmic cultures at 24h. As before, cyto-sfGFP was visible in the periplasmic fraction, suggesting a negative impact on cell integrity.

Fluorescence microscopy (Fig. 5) showed that green fluorescence was homogenous throughout bacteria expressing cyto-sfGFP and PelB^SP^::sfGFP, and punctate green fluorescence was visible for some cells expressing DsbA^sp^::sfGFP, as with the *T7lac* promoter. Once again, “halos” signifying periplasmic export were not visible. This experiment was repeated using structured illumination microscopy (SIM) which has a higher resolution (Fig. 6). Again, cyto-sfGFP and PelB^sp^::sfGFP appeared to have uniform fluorescence distribution throughout the bacteria, commensurate with cytoplasmic localisation. Cells expressing DsbA^sp^::sfGFP displayed the characteristic punctate fluorescence, although some also showed a “halo” of green fluorescence (arrowed); Co-staining the membranes with Nile Red [42] revealed that many points of green fluorescence appeared to colocalise with the envelope. This could suggest that the green “points” are periplasmic inclusion bodies. Further work is required to investigate this observation. Taken together, this data indicates that periplasmic sfGFP is not evenly distributed within the cell, leads to stress and cell death, and as such is a poor screening tool for recombinant protein production in the periplasm.

**Figure 5.**
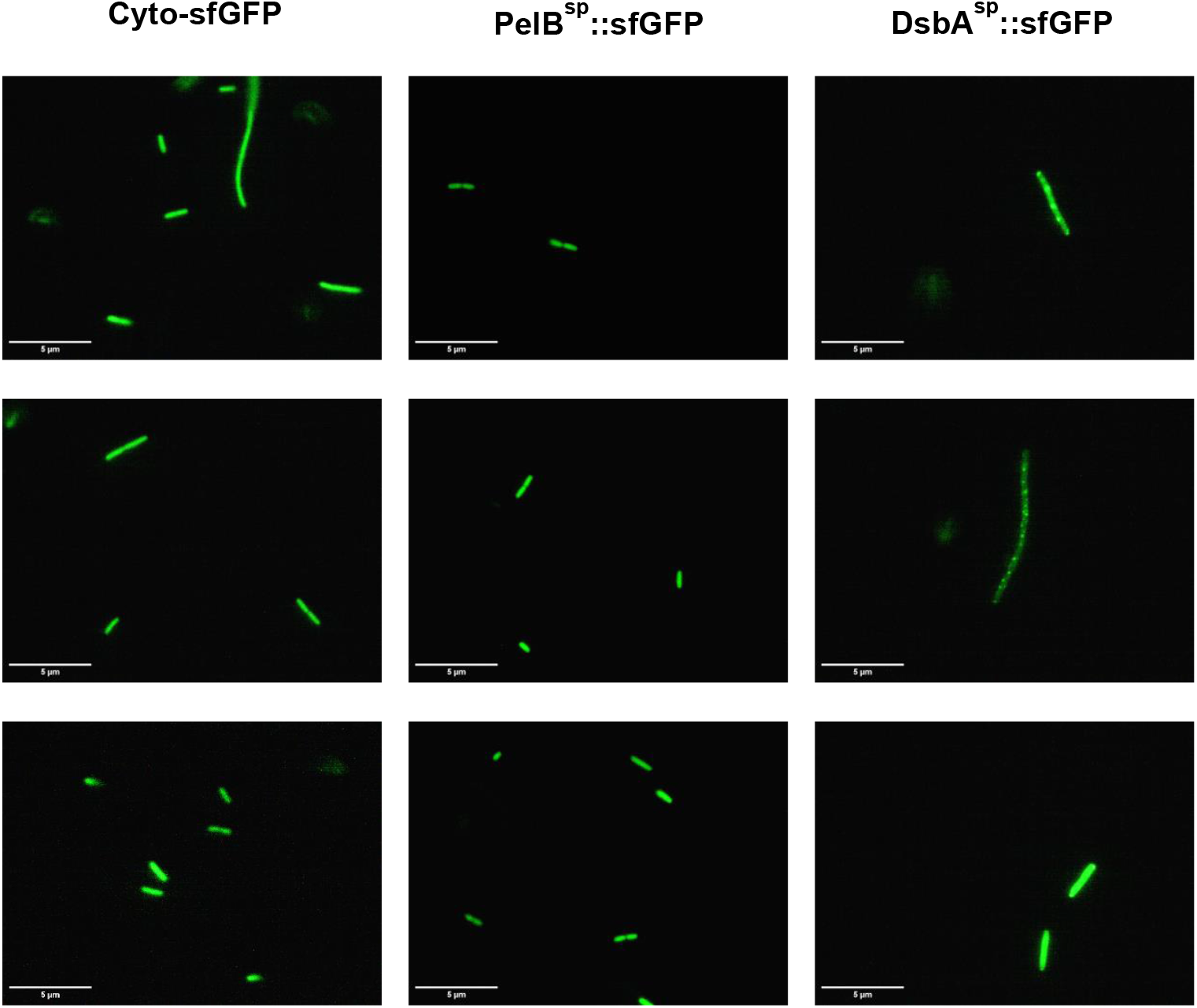
Fluorescence microscopy of sfGFP expressed from the *pBAD* promoter. *E. coli* BL21 transformed with either pBAD30-*kan*^*R*^-pelB^SP^::msfgfp, pBAD30-kan^R^-dsbA^SP^::msfgfp, or pBAD30-kan^R^-msfgfp were grown in terrific broth at 30 °C with shaking at 200 rpm until an OD_600_ ∼ 0.4-0.6, whereupon 0.2 % arabinose was added to induce RPP. At 4 h post-induction cells were visualised using fluorescence microscopy.

**Figure 6.**
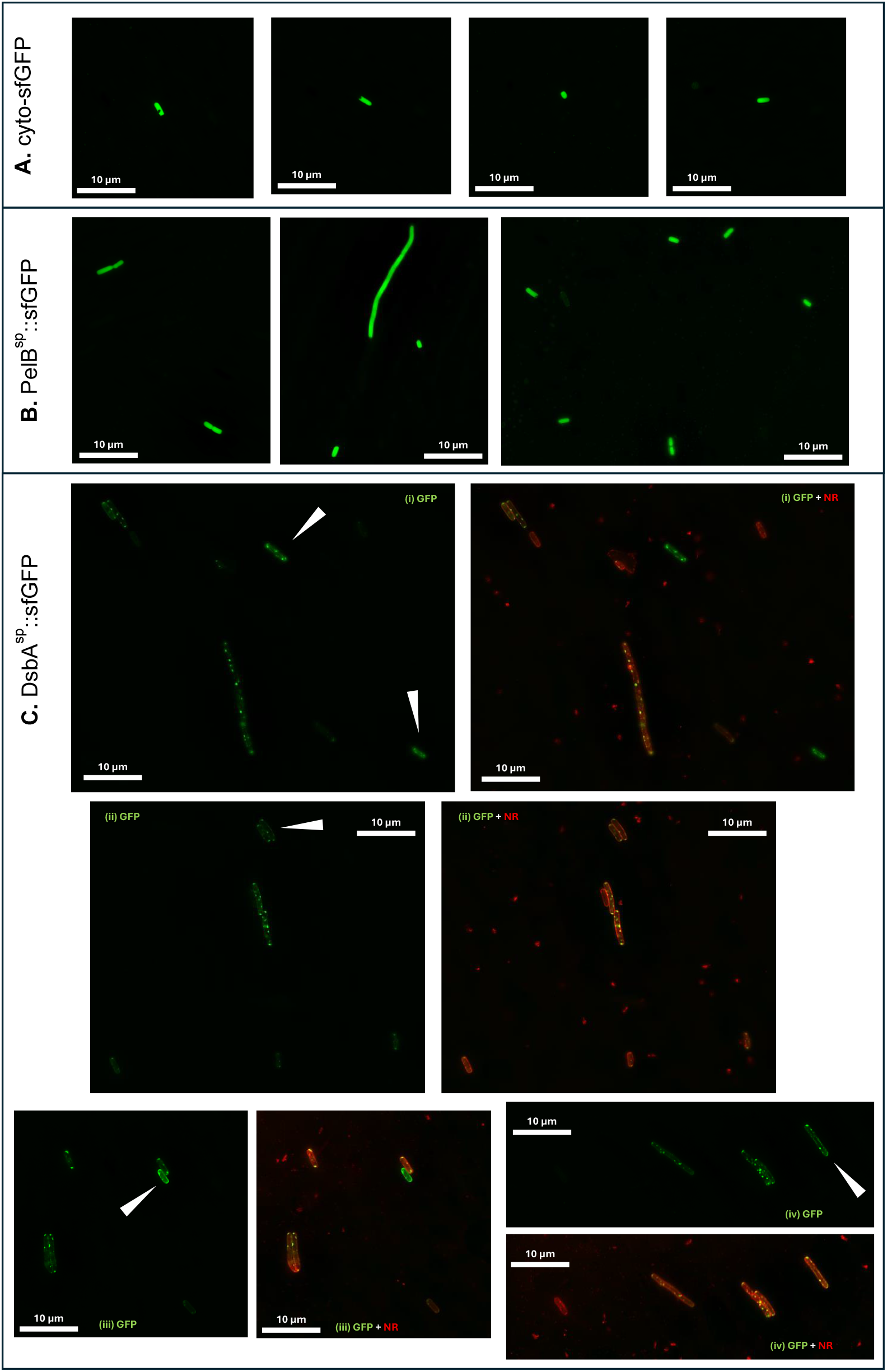
Super-resolution microscopy of sfGFP expressed from the *pBAD* promoter. *E. coli* BL21 transformed with either pBAD30-kan^R^-pelB^SP^::msfgfp, pBAD30-kan^R^-dsbA^SP^::msfgfp, or pBAD30-kan^R^-msfgfp were grown in terrific broth at 30 °C with shaking at 200 rpm until an OD_600_ ∼ 0.4-0.6, whereupon 0.2 % arabinose was added to induce RPP. At 4 h post-induction cells were visualised using SIM microscopy. Cultures expressing DsbA^sp^::sfGFP were co-stained with Nile Red.

### Cysteine-free GFP

The main theory as to why GFP presents problems for periplasmic localisation is that cysteine residues can generate errant disulphide bonds [22,23]. Although superfolder GFP has previously been reported to be functional in the periplasm [29,33] we have shown above that sfGFP gives rise to physiological stress and so is of limited use as a screening tool for periplasmic RPP. Therefore, we used a cysteine-free derivative of GFP, cfSGFP2, which carries the C48S C70M mutations and has been shown to fluoresce in the oxidizing endoplasmic reticulum in eukaryotic cells [43]. Rather than a derivative of sfGFP, cfSGFP2 is evolved from SGFP2, itself derived from EGFP [44]. SGFP2 has previously been shown to fluoresce more strongly in *E. coli* than EGFP [44] although to the best of our knowledge cfGSFP2 has not been used in *E. coli* before.

Plasmids were constructed expressing cfSGFP2 from the pBAD promoter, either without a signal peptide or with DsbA^sp^. We repeatedly attempted to synthesise a construct expressing PelB^sp^::cfSGFP2 but each time, mutations were incorporated into the resultant plasmid (in the promoter region or insertion of stop codons); we conclude that expression of PelB^sp^::cfSGFP2 is very poorly tolerated by *E. coli*, to the extent of preventing construction of an expression construct, even though pBAD30 is a low copy number “tight” expression vector without excessive leakiness.

Comparison of growth of *E. coli* BL21 expressing cytoplasmic (cyto-) cfSGFP2 and DsbA^sp^:: cfSGFP2 (Fig. 7A) reveals that cyto-cfSGFP2 does not negatively impact on growth, DsbA^sp^:: cfSGFP2 expression results in a large growth defect. Green fluorescence measurements (Fig. 7B) show that, whereas induction with a higher arabinose concentration results in higher cyto-cfSGFP2 fluorescence, fluorescence values are lower than those of sfGFP (cf Fig. 4E). DsbA^sp^::cfSGFP2 expression resulted in very low green fluorescence and cell death comparable to DsbA^sp^::sfGFP as detected by PI staining (Fig. 7C). Single-cell analysis revealed homogenous populations in respect to green fluorescence (Supplemental Fig. S5). Taken together, we conclude that periplasmic targeting of cfSGFP2 also has a negative impact on the physiology of *E. coli*, meaning that it is not a good screening tool. This also calls into question the hypothesis that errant disulphide bonding is the reason why periplasmic GFP is problematic in *E. coli*, given that cfSGFP2 does not form disulphide bonds.

**Figure 7.**
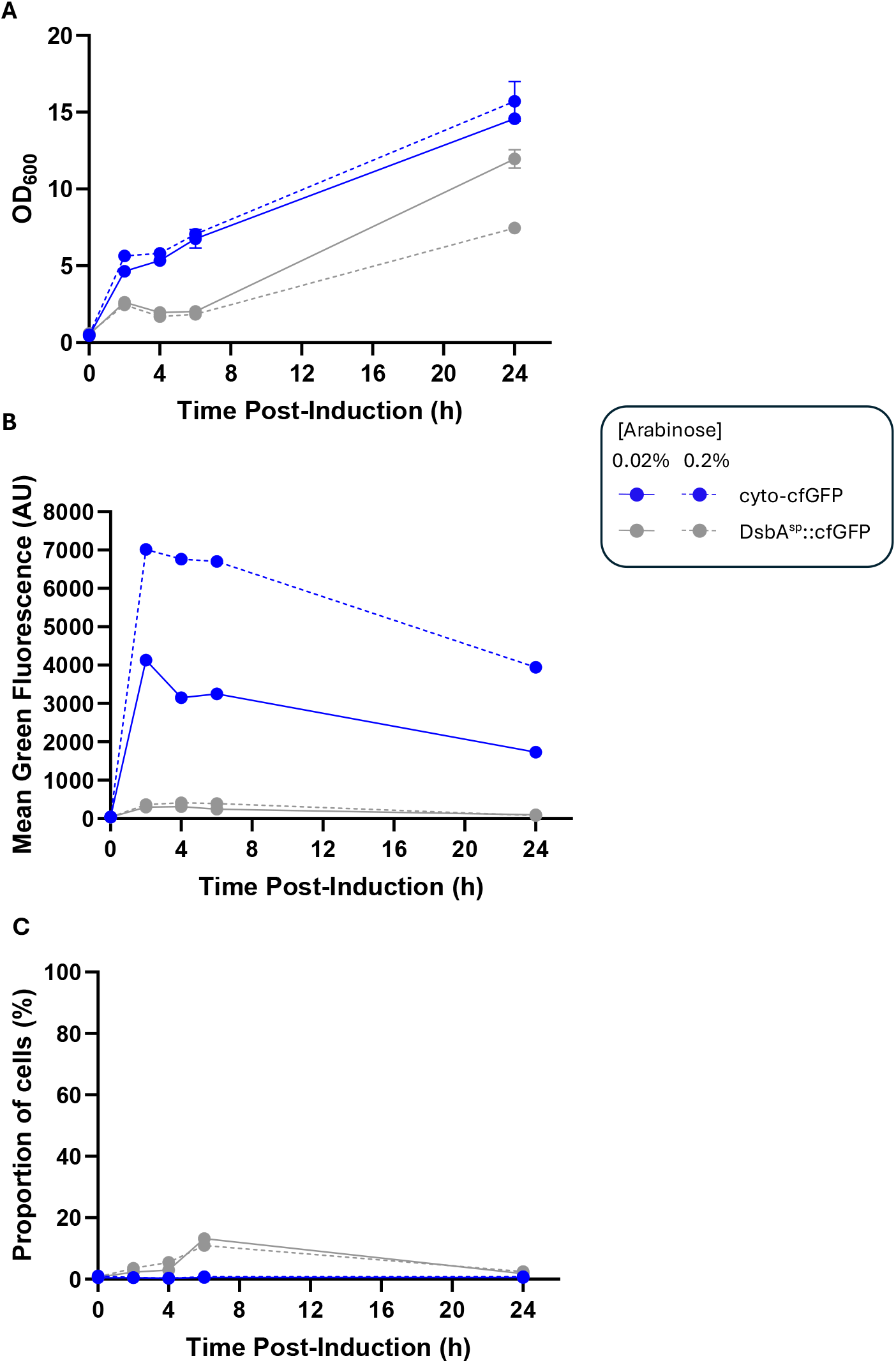
Expression of cysteine-free GFP. *E. coli* BL21 transformed with either pBAD30-kan^R^-cfsgfp2 or pBAD30-kan^R^-dsbA^SP^::cfsgfp2 were grown in terrific broth at 30 °C with shaking at 200 rpm until an OD_600_ ∼ 0.4-0.6, whereupon 0.2 % or 0.02 % arabinose was added to induce RPP. Samples were taken at regular intervals and OD_600_ was measured (**A**). Flow cytometry was used to measure green fluorescence (**B**), and the percentage of cells that were dead (PI^+^, **C**). Error bars represent +/- SD from duplicate flasks. Note that the same flow cytometer was used to generate data in Fig 7 and Fig 4D-F, so absolute fluorescence values are comparable.

## Conclusions

Since there is no way to predict which signal peptide will be optimal for each recombinant protein, and process development must determine the optimal conditions (eg growth medium, inducer concentration, growth temperature, harvest point) for each signal peptide-recombinant protein combination, screening approaches are required to accelerate process development [21]. Cytoplasmic recombinant protein production can be optimised using GFP as a fusion protein, but periplasmic translocation of GFP is problematic, even with superfolder GFP, previously reported to fold in the periplasm [29,33]. We have revealed that sfGFP poses significant physiological stress to *E. coli*, limiting growth and recombinant protein production and translocation, and thereby prevents its use as a routine screening tool. Furthermore, a proposed reason that GFP derivatives are ineffective in the periplasm, errant disulphide bonding, was tested with the use of a cysteine-free GFP, which likewise was poorly tolerated by *E. coli*. Future work could include fusions to alternative fluorescent proteins, for example mCherry and derivatives, which have been shown to be periplasmically active [29], although the impact on physiology of their periplasmic expression is not well understood.

## Abbreviations

GFP: Green Fluorescent Protein
RPP: recombinant protein production
scFv: Single chain variable fragment
sfGFP: superfolder Green Fluorescent Protein
Tat: Twin arginine translocation

## Acknowledgements

AO is funded by a UK Biotechnology and Biological Sciences Research Council PhD studentship as part of the Midlands Integrative Biosciences Training Partnership (MIBTP). The SIM was provided by the COMPARE Advanced Imaging Facility. We thank Joao Correia and Alessandro Di Maio for their support and assistance in developing and completing these experiments.

## Author contributions: CRediT

**Alexander Osgerby:** Conceptualization, Formal analysis, Investigation, Methodology, Writing – review and editing. **Tim Overton:** Conceptualization, Funding acquisition, Supervision, Writing – original draft.

## References

[1] Overton TW. Recombinant protein production in bacterial hosts. Drug Discov Today 2014;19:590–601. 10.1016/j.drudis.2013.11.008.

[2] Walsh G, Walsh E. Biopharmaceutical benchmarks 2022. Nat Biotechnol 2022;40. 10.1038/s41587-022-01582-x.

[3] Sevastsyanovich Y, Alfasi S, Griffiths L, Overton T, Cole J. Improving yield and quality of recombinant proteins in Escherichia coli by stress minimisation and mutant selection. N Biotechnol 2009;25:S244. 10.1016/j.nbt.2009.06.240.

[4] Wyre C, Overton TW. Use of a stress-minimisation paradigm in high cell density fed-batch Escherichia coli fermentations to optimise recombinant protein production. J Ind Microbiol Biotechnol 2014;41:1391–404. 10.1007/s10295-014-1489-1.

[5] Vera A, González-Montalbán N, Arís A, Villaverde A. The conformational quality of insoluble recombinant proteins is enhanced at low growth temperatures. Biotechnol Bioeng 2007;96:1101–6. 10.1002/bit.21218.

[6] Zulkifly NAH, Selas Castiñeiras T, Overton TW. Optimisation of recombinant TNFα production in Escherichia coli using GFP fusions and flow cytometry. Front Bioeng Biotechnol 2023;11. 10.3389/fbioe.2023.1171823.

[7] Hothersall J, Lai S, Zhang N, Godfrey RE, Ruanto P, Bischoff S, et al. Inexpensive protein overexpression driven by the NarL transcription activator protein. Biotechnol Bioeng 2022;119:1614–23. 10.1002/bit.28071.

[8] Hothersall J, Godfrey RE, Fanitsios C, Overton TW, Busby SJW, Browning DF. The PAR promoter expression system: Modified lac promoters for controlled recombinant protein production in Escherichia coli. N Biotechnol 2021;64:1–8. 10.1016/j.nbt.2021.05.001.

[9] Hothersall J, Osgerby A, Godfrey RE, Overton TW, Busby SJW, Browning DF. New vectors for urea-inducible recombinant protein production. N Biotechnol 2022;72:89–96. 10.1016/j.nbt.2022.10.003.

[10] Gupta SK, Shukla P. Microbial platform technology for recombinant antibody fragment production: A review. Crit Rev Microbiol 2017;43:31–42. 10.3109/1040841X.2016.1150959,.

[11] Goel N, Stephens S. Certolizumab pegol. MAbs 2010;2:137–47. 10.4161/MABS.2.2.11271,.

[12] Bardwell JCA, McGovern K, Beckwith J. Identification of a protein required for disulfide bond formation in vivo. Cell 1991;67:581–9. 10.1016/0092-8674(91)90532-4.

[13] Neu HC, Heppel LA. The release of enzymes from Escherichia coli by osmotic shock and during the formation of spheroplasts. J Biol Chem 1965;240:3685–92. 10.1016/s0021-9258(18)97200-5.

[14] Frain KM, Robinson C, van Dijl JM. Transport of Folded Proteins by the Tat System. The Protein Journal 2019 38:4 2019;38:377–88. 10.1007/S10930-019-09859-Y.

[15] Palmer T, Berks BC. The twin-arginine translocation (Tat) protein export pathway. Nat Rev Microbiol 2012;10. 10.1038/nrmicro2814.

[16] Tsirigotaki A, De Geyter J, Šoštarić N, Economou A, Karamanou S. Protein export through the bacterial Sec pathway. Nat Rev Microbiol 2017;15:21–36. 10.1038/nrmicro.2016.161.

[17] Cranford-Smith T, Huber D. The way is the goal: how SecA transports proteins across the cytoplasmic membrane in bacteria. FEMS Microbiol Lett 2018;365:93. 10.1093/FEMSLE/FNY093.

[18] Saraogi I, Shan S ou. Co-translational protein targeting to the bacterial membrane. Biochimica et Biophysica Acta (BBA) - Molecular Cell Research 2014;1843:1433–41. 10.1016/J.BBAMCR.2013.10.013.

[19] Freudl R. Signal peptides for recombinant protein secretion in bacterial expression systems. Microb Cell Fact 2018;17:52. 10.1186/s12934-018-0901-3.

[20] Selas Castiñeiras T, Williams SG, Hitchcock A, Cole JA, Smith DC, Overton TW. Development of a generic β-lactamase screening system for improved signal peptides for periplasmic targeting of recombinant proteins in Escherichia coli. Sci Rep 2018;8:6986. 10.1038/s41598-018-25192-3.

[21] Osgerby A, Overton TW. Approaches for high-throughput quantification of periplasmic recombinant proteins. N Biotechnol 2023;77. 10.1016/j.nbt.2023.09.003.

[22] Dammeyer T, Tinnefeld P. Engineered fluorescent proteins illuminate the bacterial periplasm. Comput Struct Biotechnol J 2012;3:e201210013. 10.5936/CSBJ.201210013.

[23] Jain RK, Joyce PBM, Molinete M, Halban PA, Gorr SU. Oligomerization of green fluorescent protein in the secretory pathway of endocrine cells. Biochem J 2001;360:645–9. 10.1042/0264-6021:3600645.

[24] Thomas JD, Daniel RA, Errington J, Robinson C. Export of active green fluorescent protein to the periplasm by the twin-arginine translocase (Tat) pathway in Escherichia coli. Mol Microbiol 2001;39. 10.1046/j.1365-2958.2001.02253.x.

[25] Pédelacq J-D, Cabantous S, Tran T, Terwilliger TC, Waldo GS. Engineering and characterization of a superfolder green fluorescent protein 2006. 10.1038/nbt1172.

[26] Waldo GS, Standish BM, Berendzen J, Terwilliger TC. Rapid protein-folding assay using green fluorescent protein. Nat Biotechnol 1999;17:691–5. 10.1038/10904.

[27] Crameri A, Whitehorn EA, Tate E, Stemmer WPC. Improved Green Fluorescent Protein by Molecular Evolution Using DNA Shuffling. Nature Biotechnology 1996 14:3 1996;14:315–9. 10.1038/nbt0396-315.

[28] Cormack BP, Valdivia RH, Falkow S. FACS-optimized mutants of the green fluorescent protein (GFP). Gene 1996;173:33–8. 10.1016/0378-1119(95)00685-0.

[29] Dinh T, Bernhardt TG. Using superfolder green fluorescent protein for periplasmic protein localization studies. J Bacteriol 2011;193:4984–7. 10.1128/JB.00315-11.

[30] Schierle CF, Berkmen M, Huber D, Kumamoto C, Boyd D, Beckwith J. The DsbA signal sequence directs efficient, cotranslational export of passenger proteins to the Escherichia coli periplasm via the signal recognition particle pathway. J Bacteriol 2003;185:5706–13. 10.1128/JB.185.19.5706-5713.2003.

[31] Schibich D, Gloge F, Pöhner I, Björkholm P, Wade RC, Von Heijne G, et al. Global profiling of SRP interaction with nascent polypeptides. Nature 2016 536:7615 2016;536:219–23. 10.1038/nature19070.

[32] Fisher AC, DeLisa MP. Laboratory evolution of fast-folding green fluorescent protein using secretory pathway quality control. PLoS One 2008;3. 10.1371/JOURNAL.PONE.0002351.

[33] Aronson DE, Costantini LM, Snapp EL. Superfolder GFP Is Fluorescent in Oxidizing Environments When Targeted via the Sec Translocon. Traffic 2011;12:543–8. 10.1111/J.1600-0854.2011.01168.X.

[34] Lee HC, Bernstein HD. The targeting pathway of Escherichia coli presecretory and integral membrane proteins is specified by the hydrophobicity of the targeting signal. Proc Natl Acad Sci U S A 2001;98:3471–6. 10.1073/PNAS.051484198.

[35] Rajacharya GH, Kumar J, Gupta JA, Rathore AS. Impact of stringent stress response and amino acid supplementation on recombinant protein production in Escherichia coli. Biochem Eng J 2025;219:109727. 10.1016/J.BEJ.2025.109727.

[36] de Marco A. Recent advances in recombinant production of soluble proteins in E. coli. Microb Cell Fact 2025;24:21. 10.1186/S12934-025-02646-8.

[37] Studier FW, Moffatt BA. Use of bacteriophage T7 RNA polymerase to direct selective high-level expression of cloned genes. J Mol Biol 1986;189:113–30. 10.1016/0022-2836(86)90385-2.

[38] Anton BP, Raleigh EA. Complete Genome Sequence of NEB 5-alpha, a Derivative of Escherichia coli K-12 DH5α. Genome Announc 2016;4. 10.1128/GENOMEA.01245-16.

[39] Skinner SO, Sepúlveda LA, Xu H, Golding I. Measuring mRNA copy number in individual Escherichia coli cells using single-molecule fluorescent in situ hybridization. Nature Protocols 2013 8:6 2013;8:1100–13. 10.1038/nprot.2013.066.

[40] Schneider CA, Rasband WS, Eliceiri KW. NIH Image to ImageJ: 25 years of image analysis. Nat Methods 2012;9:671–5. 10.1038/nmeth.2089.

[41] Guzman LM, Belin D, Carson MJ, Beckwith J. Tight regulation, modulation, and high-level expression by vectors containing the arabinose P(BAD) promoter. J Bacteriol 1995;177:4121–30. 10.1128/jb.177.14.4121-4130.1995.

[42] Spahn CK, Glaesmann M, Grimm JB, Ayala AX, Lavis LD, Heilemann M. A toolbox for multiplexed super-resolution imaging of the E. coli nucleoid and membrane using novel PAINT labels. Scientific Reports 2018 8:1 2018;8:1–12. 10.1038/s41598-018-33052-3.

[43] Suzuki T, Arai S, Takeuchi M, Sakurai C, Ebana H, Higashi T, et al. Development of Cysteine-Free Fluorescent Proteins for the Oxidative Environment. PLoS One 2012;7:e37551. 10.1371/JOURNAL.PONE.0037551.

[44] Kremers GJ, Goedhart J, Van Den Heuvel DJ, Gerritsen HC, Gadella TWJ. Improved green and blue fluorescent proteins for expression in bacteria and mammalian cells. Biochemistry 2007;46:3775–83. 10.1021/BI0622874/SUPPL_FILE/BI0622874SI20061105_060255.PDF.

