## Supplemental information for "Periplasmic production of Green Fluorescent Protein is poorly tolerated by *Escherichia coli*"

### SUPPLEMENTAL FIGURES

**Figure S1**

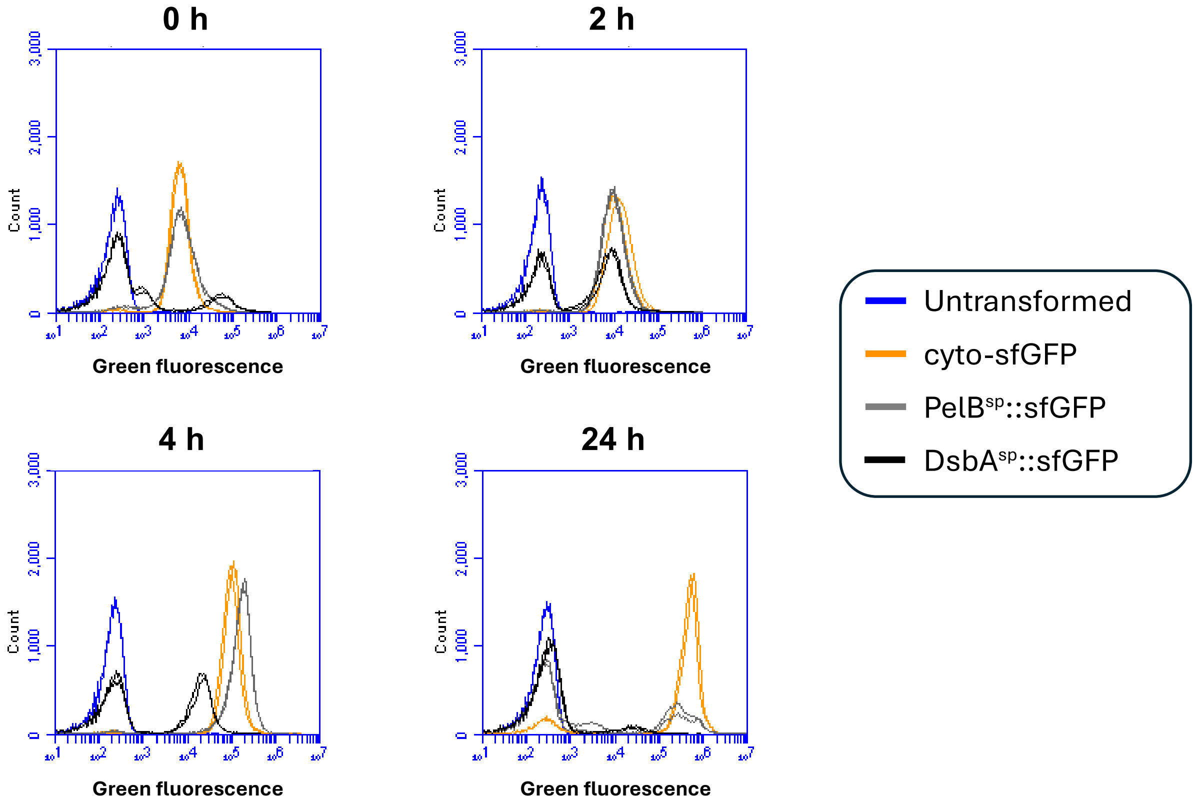


*E. coli* BL21(DE3) transformed with either pET-26b(+)Δ-ATG-msfgfp (expressing cyto-sfGFP), pET-26b(+)-PelB$^{SP}$::msfgfp (periplasmic targeting using PelB$^{sp}$) or pET-26b(+)Δ-DsbA$^{SP}$::msfgfp (periplasmic targeting using DsbA$^{sp}$), and an untransformed control were grown in terrific broth at 30 °C with shaking at 200 rpm until an $OD_{600}$ ~ 0.4-0.6, whereupon 50 μM IPTG was added. Flow cytometry was used to measure green fluorescence; each condition is shown in duplicate.

**Figure S2**

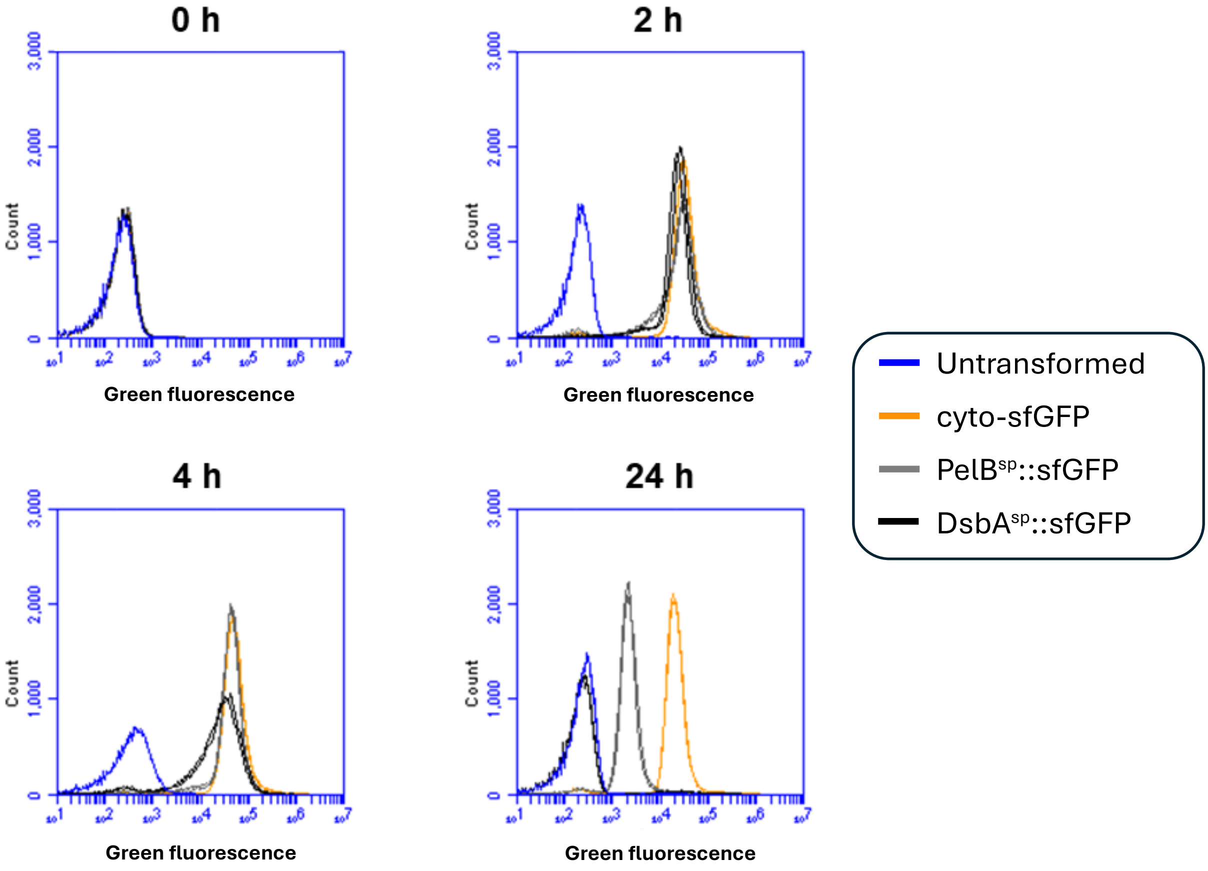


*E. coli* BL21 transformed with either pBAD30-kan$^{R}$-pelB$^{SP}$::msfgfp, pBAD30-kan$^{R}$-dsbA$^{SP}$::msfgfp, or pBAD30-kan$^{R}$-msfgfp (and an untransformed control) were grown in terrific broth at 30 °C with shaking at 200 rpm until an $OD_{600}$ ~ 0.4-0.6, whereupon 0.2 % arabinose was added to induce RPP. Samples were taken at regular intervals and flow cytometry was used to measure green fluorescence. Each condition is shown in duplicate.

**Figure S3**

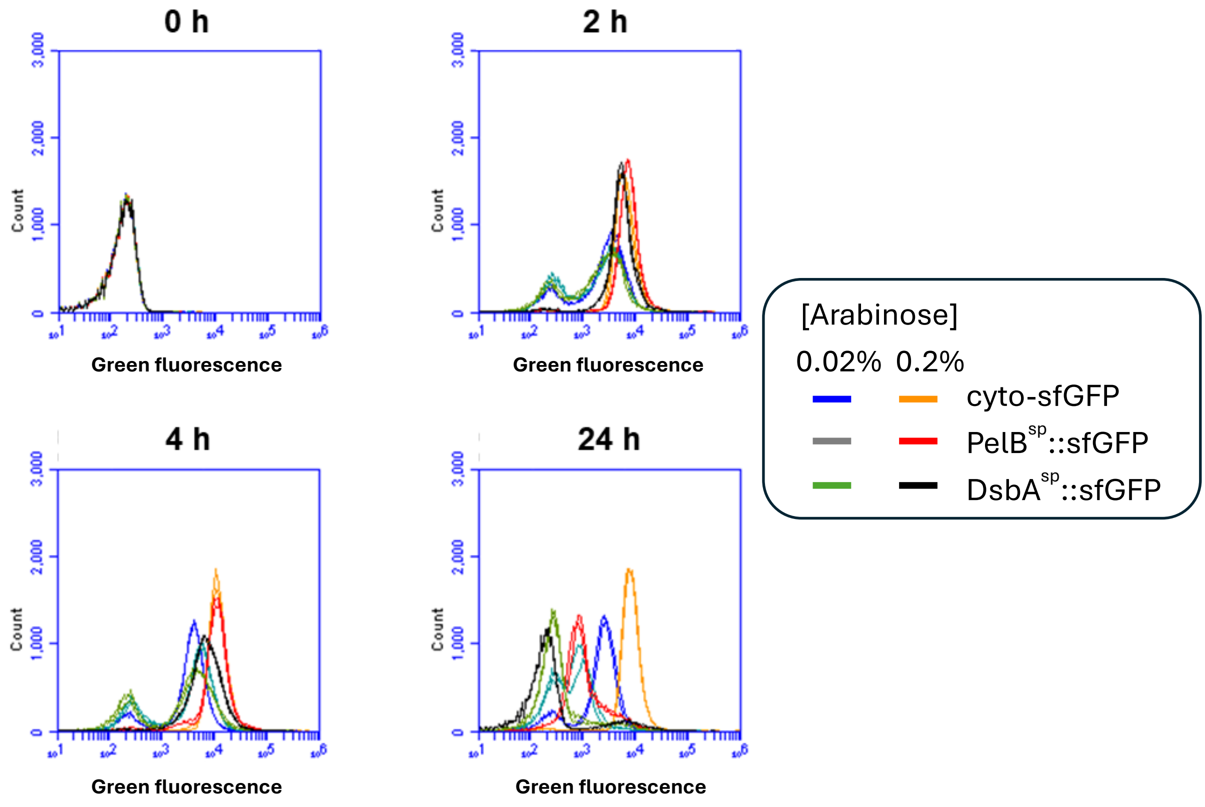


*E. coli* BL21 transformed with either pBAD30-kan$^R$-pelB$^{SP}$::msfgfp, pBAD30-kan$^R$-dsbA$^{SP}$::msfgfp, or pBAD30-kan$^R$-msfgfp were grown in terrific broth at 30 °C with shaking at 200 rpm until an $OD_{600}$ ~ 0.4-0.6, whereupon 0.02 % or 0.2 % arabinose was added to induce RPP. Samples were taken at regular intervals and flow cytometry was used to measure green fluorescence. Each condition is shown in duplicate.

**Figure S4**

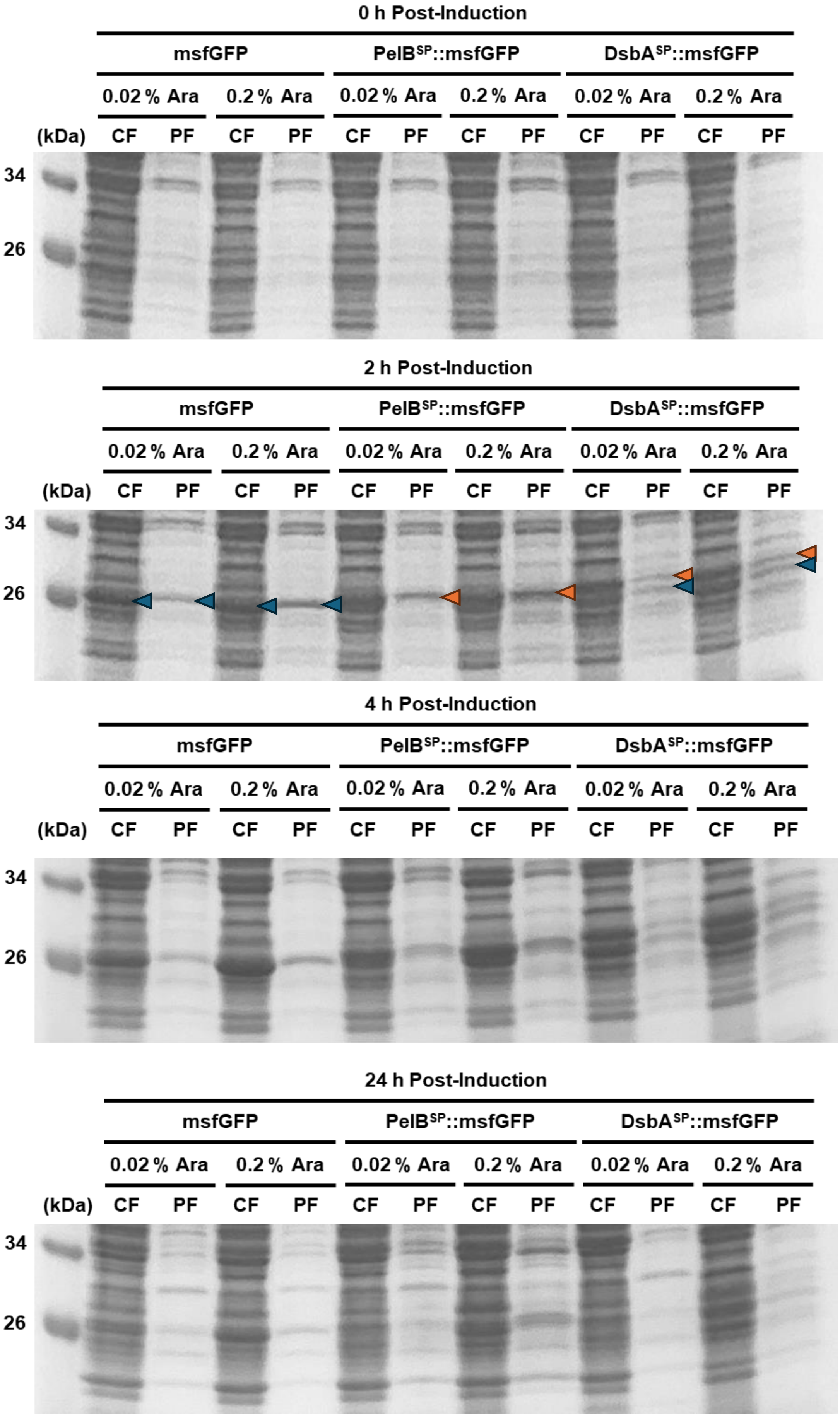
0 h Post-Induction
msfGFP
PelB$^{SP}$::msfGFP
DsbA$^{SP}$::msfGFP
0.02 % Ara
0.2 % Ara
(kDa)
CF
PF
34
26
2 h Post-Induction
msfGFP
PelB$^{SP}$::msfGFP
DsbA$^{SP}$::msfGFP
0.02 % Ara
0.2 % Ara
(kDa)
CF
PF
34
26
4 h Post-Induction
msfGFP
PelB$^{SP}$::msfGFP
DsbA$^{SP}$::msfGFP
0.02 % Ara
0.2 % Ara
(kDa)
CF
PF
34
26
24 h Post-Induction
msfGFP
PelB$^{SP}$::msfGFP
DsbA$^{SP}$::msfGFP
0.02 % Ara
0.2 % Ara
(kDa)
CF
PF
34
26

*E. coli* BL21 transformed with either pBAD30-kan$^{R}$-pelB$^{SP}$::msfgfp, pBAD30-kan$^{R}$-dsbA$^{SP}$::msfgfp, or pBAD30-kan$^{R}$-msfgfp were grown in terrific broth at 30 °C with shaking at 200 rpm until an $OD_{600}$ ~ 0.4-0.6, whereupon 0.02 % or 0.2 % arabinose was added to induce RPP. Samples were taken at regular intervals, fractionated into periplasmic (PF) and cytoplasmic (CF) fractions, and separated by SDS-PAGE. Bands corresponding to sfGFP are indicated by arrows on the 2 h gel, blue referring to sfGFP and red to sfGFP fused to a signal peptide with a resultant higher molecular weight.

**Figure S5**

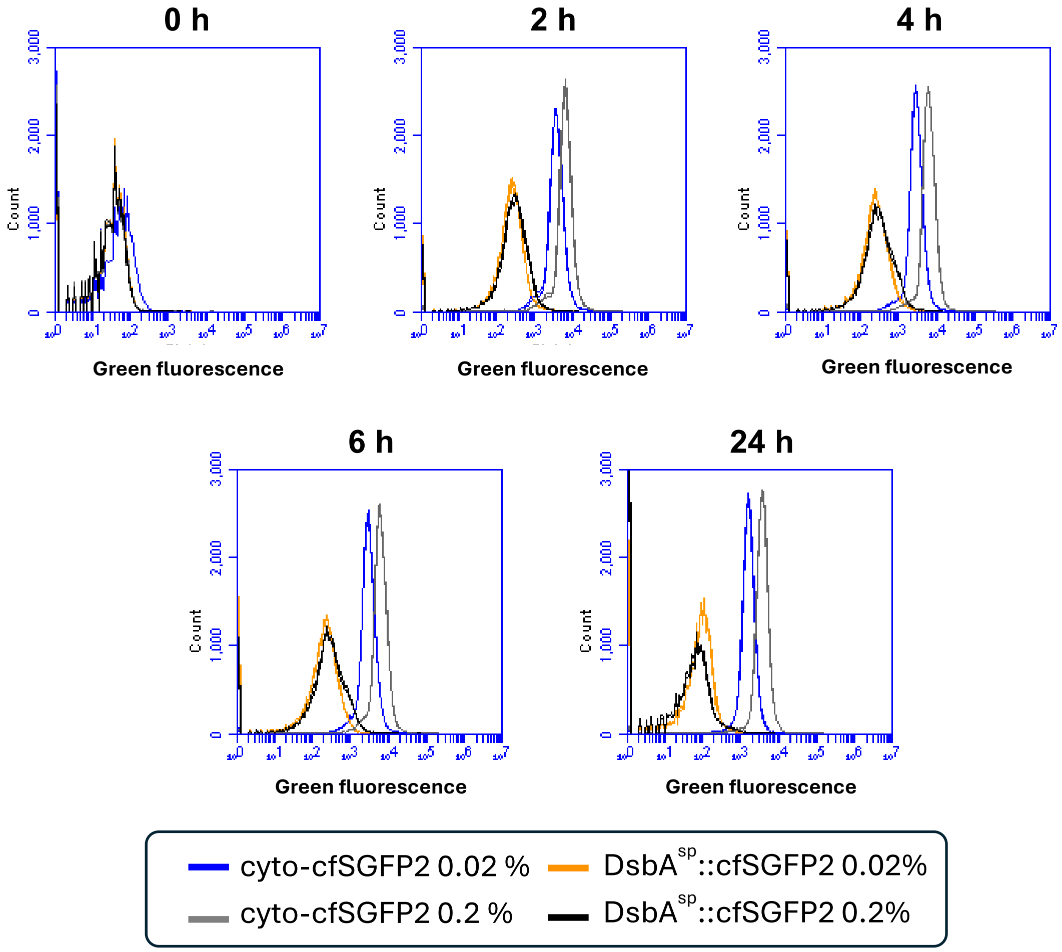


*E. coli* BL21 transformed with either pBAD30-kan$^{R}$-cfsgfp2 or pBAD30-kan$^{R}$-dsbA$^{SP}$::cfsgfp2 were grown in terrific broth at 30 °C with shaking at 200 rpm until an $OD_{600}$ ~ 0.4-0.6, whereupon 0.2 % or 0.02 % arabinose was added to induce RPP. Samples were taken at regular intervals and flow cytometry was used to measure green fluorescence. Each condition is shown in duplicate.

**Supplementary table S1.** Plasmids used in this work

| Plasmid | Details | Source |
|---|---|---|
| pET-26b(+) | Expression vector, T7*lac* promoter, *lacI* and PelB$^{SP}$ (Kan$^{R}$). | Kindly provided by Dr Douglas Browning (Novagen). |
| pET GFP LIC (u-msfgfp) | Cloning plasmid containing msfGFP (Amp$^{R}$). | Kindly provided by Prof Jessica Blair (Addgene plasmid #29772). |
| pET-26b(+)-pelB$^{SP}$::msfgfp | Variant of pET-26b(+) with *msfgfp* inserted downstream of *pelB*$^{SP}$ (Kan$^{R}$). | This work. |
| pET-26b(+)Δ-dsbA$^{SP}$::msfgfp | Variant of pET-26b(+) with *pelB*$^{SP}$ deleted and replaced with *dsbA*$^{SP}$ and *msfgfp* inserted downstream (Kan$^{R}$). | This work. |
| pET-26b(+)Δ-GTG-msfgfp | Variant of pET-26b(+) with *pelB*$^{SP}$ deleted and *msfgfp* inserted downstream with a GTG start codon (Kan$^{R}$). | This work. |
| pET-26b(+)Δ-ATG-msfgfp | Variant of pET-26b(+)Δ-*msfgfp* with the GTG start codon replaced with the canonical ATG (Kan$^{R}$). | This work. |
| pBAD30 | Empty expression plasmid expressing AraC and with $P_{araBAD}$ ustream of MCS (Amp$^{R}$). | Kindly provided by Professor Steve Busby (Guzman et al., 1995). |
| pBAD30-kan$^{R}$ | Variant of pBAD30 with *amp*$^{R}$ replaced with *kan*$^{R}$ and *rrnB* terminators in the opposing orientation (Kan$^{R}$). | This work. |
| pBAD30-kan$^{R}$-pelB$^{SP}$::msfgfp | Variant of pBAD30-*kan*$^{R}$ with *pelB*$^{SP}$::*msfgfp* inserted downstream of $P_{araBAD}$ (Kan$^{R}$). | This work. |
| pBAD30-kan$^{R}$-dsbA$^{SP}$::msfgfp | Variant of pBAD30-*kan*$^{R}$ with *dsbA*$^{SP}$::*msfgfp* inserted downstream of $P_{araBAD}$ (Kan$^{R}$). | This work. |
| pBAD30-kan$^{R}$-msfgfp | Variant of pBAD30-*kan*$^{R}$ with *msfgfp* inserted downstream of $P_{araBAD}$ (Kan$^{R}$). | This work. |
| pBAD30-kan$^{R}$-cfsgfp2 | Variant of pBAD30-*kan*$^{R}$ with *cfsgfp2* inserted downstream of $P_{araBAD}$ (Kan$^{R}$). | This work. |
| pBAD30-kan$^{R}$-dsbA$^{SP}$::cfsgfp2 | Variant of pBAD30-*kan*$^{R}$ with *dsbA*$^{SP}$::*cfsgfp2* inserted downstream of $P_{araBAD}$ (Kan$^{R}$). | This work. |

**Supplementary Table S2.** Oligonucleotide primers used in this work

| Primer Name | Sequence (5'-3') |
|---|---|
| pET26b_Fwd_GA_1 | GACGAGCTGTACAAGTGAGATCCGGCTGCTAACAAAGCCCG |
| pET26b_Rev_GA_1 | CTCCTCGCCCTTGCTGGCCATCGCCGGCTGGG |
| msfGFP_Fwd_GA_1 | CAGCCGGCGATGGCCAGCAAGGGCGAGGAGCTG |
| msfGFP_Rev_GA_1 | AGCAGCCGGATCTCACTTGTACAGCTCGTCCATGCC |
| pET26b_Fwd_GA_3 | TAGTAACCAACTCCATGCTAACAAAGCCCGAAA |
| pET26b_Rev_GA_3 | CCAAATCTTTTTCATATGTATATCTCCTTCTTAAAGTTAAAC |
| DsbA-msfGFP_Fwd_5 | TGAAAAAGATTTGGCTGGCGCTGGCTGGTTTAGTTTTAGCGTTTAGCGCATCGGCGAGCAAGGGCGAGGAGCTG |
| DsbA-msfGFP_Rev_5 | CTTCCTTTCGGGCTTTGTTAGCA |
| DsbA-msfGFP_Fwd_GA_6 | GAAGGAGATATACATATGAAAAAGATTTGGCTGGC |
| DsbA-msfGFP_Rev_GA_6 | CGGGCTTTGTTAGCATGGAGTTGGTTACTACTTGT |
| pET26b_Fwd_GA_2 | CTAACAAAGCCCGAAGCTAACAAAGCCCGAAAG |
| pET26b_Rev_GA_2 | CTCGCCCTTGCTCACATGTATATCTCCTTCTTAAAGTTAAAC |
| msfGFP_Fwd_GA_4 | GAAGGAGATATACATGTGAGCAAGGGCGAGGAG |
| msfGFP_Rev_GA_4 | TCGGGCTTTGTTAGCTTCGGGCTTTGTTAGCAGC |
| pET-msfGFP_Sub_Fwd | AGATATACATATGAGCAAGGGCG |
| pET-msfGFP_Sub_Rev | CCTTCTTAAAGTTAAACAAAATTATTTCTAG |
| pET26b_Fwd_CP_1 | GTGATGTCGGCGATATAGG |
| pET26b_Rev_CP_1 | TTCCTTTCGGGCTTTGTTAG |
| T7 Terminator | GCTAGTTATTGCTCAGCGG |
| pBAD30_Fwd_GA_1 | GATCAAAGGATCTTCCTGTCAGACCAAGTTTACTC |
| pBAD30_Rev_GA_1 | AACGTTGCGAAGCAAAGAGTTTGTAGAAACGCAAA |

| | |
|---|---|
| kan_Fwd_GA_1 | AACTTGGTCTGACAGGAAGATCCTTTGATCTTTTCTAC |
| kan_Rev_GA_1 | ACTTCTGAGTTCGGCTTAGAAAAACTCATCGAGCATC |
| rrnB T1_Fwd | GATGAGTTTTTCTAAGCCGAACTCAGAAGTGAAAC |
| rrnB T1_Rev | GTTTCTACAAACTCTTTGCTTCGCAACGTTCAAAT |
| pBAD30_Fwd_CP_1 | GGCATCAAATTAAGCAGAAGG |
| pBAD30_Rev_CP_1 | GTAAAGCACTAAATCGGAACC |
| pBAD30_Fwd_GA_2 | TTGAGGGGTTTTTTGAATCAGAACGCAGAAGCG |
| pBAD30_Rev_GA_2 | TTAAACAAAATTATTAAACGGGTATGGAGAAACAG |
| pET26b-msfGFP_Fwd_GA_1 | TCTCCATACCCGTTTAATAATTTTGTTTAACTTTAAGAAGGA |
| pET26b-msfGFP_Rev_GA_1 | TTCTGCGTTCTGATTCAAAAAACCCCTCAAGAC |
| pBAD30_Fwd_CP_2 | CTTTCATTCCCAGCGGTC |
| pBAD30_Rev_CP_3 | CGCCAGGCAAATTCTGTT |
| pBAD30-Cyto_Fwd | CCAACTCCATAAGGATCC |
| pBAD30-Cyto_Rev | ATGTATATCTCCTTCTTAAAGTTAAAC |
| pBAD30-Cyto_Fwd_GA | GAGCTGTACAAGTAACCAACTCCATAAGGATCC |
| pBAD30-Cyto_Rev_GA | GCCCTTGCTCACCATATGTATATCTCCTTCTTAAAGTTAAAC |
| cfSGFP2_Fwd_1 | TCGTGCTGTGTTAGATGGTG |
| cfSGFP2_Rev_1 | GTTCGTGAGTCGTCAGTTAC |
| cfGFP_Fwd_GA_1 | GAAGGAGATATACATATGGTGAGCAAGGGCGAG |
| cfGFP_Rev_GA_1 | TCCTTATGGAGTTGGTTACTTGTACAGCTCGTCCATGC |
| pBAD30-cfSGFP2_Fwd | GTGAGCAAGGGCGAGGAG |
| pBAD30-cfSGFP2_Rev | AAACGGGTATGGAGAAACAGTAGAG |
| pBAD30-DsbA-cfSGFP2_Fwd | ATGAAAAAGATTTGGCTGGCGCTGGCTGGTTTAGTTTTAGCGTTTAGCGCATCGGCGGTGAGCAAGGGCGAGGAG |
| pBAD30-cfSGFP2_Rev_2 | ATGTATATCTCCTTCTTAAAGTTAAACAAAATTATTAAACGGGTATGGAGAAACAGTAGAG |

**Supplemental methods – construction of plasmids**

*General cloning methods*

Plasmid constructs were generated by PCR and HiFi DNA assembly (New England Biolabs) or blunt end ligation. Following amplification, PCR products were verified by agarose gel electrophoresis. Template DNA in backbone amplicons was DpnI digested (New England Biolabs). After purification, DNA concentration of backbone and insert were quantified by Nanodrop (Thermo Scientific) and assembled by HiFi DNA assembly. Reactions were transformed into chemically competent *E. coli* DH5α cells (New England Biolabs 5-alpha) and plated on LB agar with 50 µg·mL$^{-1}$ Kanamycin. Transformants were screened by colony PCR. Positive clones were grown overnight in LB with 50 µg·mL$^{-1}$ Kanamycin and constructs purified by miniprep (QIAGEN QIAprep spin miniprep kit). Constructs were verified by Sanger sequencing (Source Bioscience, Nottingham, UK).

*pET-based plasmids*

Constructs were verified by sequencing using primer pET26b_Fwd_CP_1.

pET-26b(+)_pelB$^{SP}$::msfgfp was assembled by PCR amplifying pET-26b(+) using primers pET26b_Fwd_GA_1 and pET26b_Rev_GA_1, deleting the multiple cloning site (MCS). The *msfgfp* insert was PCR amplified from pET GFP LIC using primers msfGFP_Fwd_GA_1 and msfGFP_Rev_GA_1. Backbone and insert were purified by PCR purification and assembled by HiFi DNA assembly, fusing the *msfgfp* insert to the 3' end of *pelB*$^{SP}$. Transformants were screened by colony PCR using primers pET26b_Fwd_CP_1 and pET26b_Rev_CP_1.

pET-26b(+)_dsbA$^{SP}$::msfgfp was assembled by PCR amplifying pET-26b(+)using primers pET26b_Fwd_GA_3 and pET26b_Rev_GA_3, deleting *pelB*$^{SP}$ and the MCS. The pET-26b(+)Δ backbone was gel purified. The *dsbA*$^{sp}$::msfGFP insert was prepared in two steps. First, pET GFP LIC (u-msfGFP) was PCR amplified using primers DsbA-msfGFP_Fwd_5, which incorporated the entirety of *dsbA*$^{SP}$ at its 5' end, and DsbA-msfGFP_Rev_5. The *dsbA*$^{SP}$::*msfgfp* fusion was purified by PCR purification (QIAGEN QIAquick PCR purification kit). Primers DsbA-msfGFP_Fwd_GA_6 and DsbA-msfGFP_Rev_GA_6 were used to PCR amplify *dsbA*$^{SP}$::*msfgfp* creating the insert which was gel purified. During HiFi DNA assembly, *dsbA*$^{SP}$::*msfgfp* was inserted to replace *pelB*$^{SP}$ in the backbone. Transformants were screened by colony PCR using primers pET26b_Fwd_CP_1 and pET26b_Rev_CP_1.

pET-26b(+)Δ_GTG_msfgfp was assembled by PCR amplifying pET-26b(+) using primers pET26b_Fwd_GA_2 and pET26b_Rev_GA_2, deleting *pelB*$^{SP}$ and the MCS. The *msfgfp* insert was PCR amplified from pET GFP LIC (u-msfGFP) using primers msfGFP_Fwd_GA_4 and msfGFP_Rev_GA_4. The pET-26b(+)Δ backbone and *msfgfp* insert were gel purified and assembled by HiFi-DNA assembly, inserting *msfgfp* to replace *pelB*$^{SP}$ in the backbone. Transformants were screened by colony PCR using primers pET26b_Fwd_CP_1 and T7_Terminator.

pET-26b(+)Δ_ATG_msfgfp was PCR amplified from pET-26b(+)Δ_GTG_msfgfp using primers pET-msfGFP_Sub_Fwd and pET-msfGFP_Sub_Rev, annealing back to back. Forward primer pET-msfGFP_Sub_Fwd contained a single G>A point mutation in the middle of its sequence, mutagenising the *msfgfp* GTG start codon in pET-26b(+)Δ_GTG_msfgfp to ATG. The linear fragment following PCR was gel purified and then re-circularised by blunt-end ligation. Transformants were screened by colony PCR using primers pET26b_Fwd_CP_1 and T7_Terminator.

*pBAD with kanamycin resistance*

pBAD30-kanR was assembled by PCR amplifying pBAD30 using primers pBAD30_Fwd_GA_1 and pBAD30_Rev_GA_1 to prepare the backbone, deleting *ampR*. The kanamycin resistance gene was PCR amplified from pET-26b(+)using primers kan_Fwd_GA_1 and kan_Rev_GA_1 for the first insert. A second insert containing the *rrnB* T1 terminator was PCR amplified from pBAD30 using primers rrnB T1_Fwd and rrnB T1_Rev. Backbone and inserts were purified by PCR purification and assembled by HiFi DNA assembly. Primers for both inserts were designed so that *kanR* and *rrnB* T1 were assembled in the reverse orientation to *ampR*. Transformants were verified by sequencing using primers pBAD30_Fwd_CP_1 and pBAD30_Rev_CP_1.

pBAD30-kan$^{R}$-pelB$^{SP}$::msfgfp was assembled by PCR amplifying pBAD30-kanR using primers pBAD30_Fwd_GA_2 and pBAD30_Rev_GA_2 to prepare the backbone, deleting the MCS. For the insert, a region containing the T7 gene 10 RBS and *pelB$^{SP}$::msfgfp* was PCR amplified from pET-26b(+)-pelB$^{SP}$::msfgfp using primers pET26b-msfGFP_Fwd_GA_1 and pET26b-msfGFP_Rev_GA_1. Backbone and insert were purified by PCR purification. During HiFi DNA assembly, the T7 gene 10 RBS and *pelB$^{SP}$::msfgfp* fragment was inserted downstream of $P_{araBAD}$ in the same orientation, and upstream of the *rrnB* T1 and T2 terminators. Transformants were screened by colony PCR using primers pBAD30_Fwd_CP_2 and pBAD30_Rev_CP_3, then verified by sequencing using primers pBAD30_Fwd_CP_1.

pBAD30-kan$^{R}$-dsbA$^{SP}$::msfgfp was assembled by PCR amplifying pBAD30-kanR using primers pBAD30_Fwd_GA_2 and pBAD30_Rev_GA_2 to prepare the backbone, deleting the MCS. For the insert, a region containing the T7 gene 10 RBS and *dsbA$^{SP}$::msfgfp* was PCR amplified from pET-26b(+)Δ-dsbA$^{SP}$::msfgfp using primers pET26b-msfGFP_Fwd_GA_1 and pET26b-msfGFP_Rev_GA_1. Backbone and inserts were purified by PCR purification. During HiFi DNA assembly, the T7 gene 10 RBS and *dsbA$^{SP}$::msfgfp* fragment was inserted downstream of $P_{araBAD}$ in the same orientation, and upstream of the *rrnB* T1 and T2 terminators. Transformants were screened by colony PCR using primers pBAD30_Fwd_CP_2 and pBAD30_Rev_CP_3, then verified by sequencing using primers pBAD30_Fwd_CP_1.

pBAD30-kan$^{R}$-msfgfp was assembled by PCR amplifying pBAD30-kanR using primers pBAD30_Fwd_GA_2 and pBAD30_Rev_GA_2 to prepare the backbone, deleting the MCS. For the insert, a region containing the T7 gene 10 RBS and *msfgfp* was PCR amplified from pET-26b(+)Δ-ATG-msfgfp using primers pET26b-msfGFP_Fwd_GA_1 and pET26b-msfGFP_Rev_GA_1. Backbone and inserts were purified by PCR purification. During HiFi DNA assembly, the T7 gene 10 RBS and msfgfp fragment was inserted downstream of $P_{araBAD}$ in the same orientation, and upstream of the *rrnB* T1 and T2 terminators. Transformants were screened by colony PCR using primers pBAD30_Fwd_CP_2 and pBAD30_Rev_CP_3, then verified by sequencing using primers pBAD30_Fwd_CP_1.

*pBAD cfSGFP2 Constructs*

Backbone for assembly of pBAD30-kan$^R$-cfsgfp2 was prepared via PCR in two steps. First, msfgfp was deleted from pBAD30-kanR-msfgfp using primers pBAD30_Cyto_Fwd and pBAD30_Cyto_Rev, followed by amplification using primers pBAD30-Cyto_Fwd_GA and pBAD30-Cyto_Rev_GA. Both fragments to prepare the backbone were purified by gel purification. To prepare the insert, a fragment containing cfsgfp2 flanked by to optimal priming locations was chemically synthesised by GENEWIZ using the FragmentGENE service. First, the cfsgfp2 fragment was amplified using primers cfSGFP2_Fwd_1 and cfSGFP2_Rev_1, followed by a second amplification using primers cfSGFP2_Fwd_GA_1 and cfSGFP2_Rev_GA_1. Both fragments to prepare the insert were purified by PCR purification. During HiFi DNA assembly, *cfsgfp2* was inserted into the pBAD30-kanR backbone in the same position as *msfgfp* had been. Following transformation, clones were verified by Sanger sequencing using primers pBAD30_Fwd_CP_1.

Backbone for pBAD30-kan$^R$-dsbA$^{SP}$::cfsgfp2 was prepared via PCR in two steps. Due to differences in annealing temperatures for necessary priming locations, a small region between $P_{araBAD}$ and the start of *cfsgfp2*, which included the T7 gene 10 RBS, was deleted from pBAD30-kanR-cfsgfp2 using primers pBAD30-cfSGFP2_Fwd and pBAD30-cfSGFP2_Rev. This fragment was purified by PCR purification and a second round of amplification fused *dsbA$^{SP}$* immediately upstream of *cfsgfp2* and re-inserted the deleted region, using primers pBAD30-DsbA-cfSGFP2_Fwd and pBAD30-cfSGFP2_Rev_2. The forward primer contained the entire *dsbA$^{SP}$* coding sequence and reverse primer the previously deleted region at their 5' ends. This fragment was purified by gel purification and blunt-end ligation used to reassemble the construct. Following transformation, clones were verified by Sanger sequencing using primers pBAD30_Fwd_CP_1.
